# Genetic dissection of cyclic di-GMP signaling in *Pseudomonas aeruginosa via* diguanylate cyclase disruption

**DOI:** 10.1101/2025.01.26.634905

**Authors:** Román A. Martino, Daniel C. Volke, Albano H. Tenaglia, Paula M. Tribelli, Pablo I. Nikel, Andrea M. Smania

**Author notes:** Correspondence to:* **Andrea M. Smania;** Centro de Investigaciones en Química Biológica de Córdoba (CIQUIBIC-CONICET), Universidad Nacional de Córdoba, X5000HUA Córdoba, Argentina; *E-mail:*, **Pablo I. Nikel,** The Novo Nordisk Foundation Center for Biosustainability, Technical University of Denmark, 2800 Kongens Lyngby, Denmark; *E-mail:. These authors contributed equally to this work (Román A. Martino, Daniel C. Volke).

## Abstract

The second messenger *bis*-(3′→5′)-cyclic dimeric guanosine monophosphate (c-di-GMP) governs adaptive responses in the opportunistic pathogen *Pseudomonas aeruginosa*, including biofilm formation and the transition from acute to chronic infections. Understanding the intricate c-di-GMP signaling network remains challenging due to the overlapping activities of numerous diguanylate cyclases (DGCs). In this study, we employed a CRISPR-based multiplex genome-editing tool to disrupt all 32 GGDEF domain-containing proteins (GCPs) implicated in c-di-GMP signaling in *P*. *aeruginosa* UCBPP-PA14. Phenotypic and physiological analyses revealed that the resulting mutant was unable to form biofilms and had attenuated virulence. Residual c-di-GMP levels were still detected despite the extensive GCP disruption, underscoring the robustness of this regulatory network. Taken together, these findings provide insights into the complex c-di-GMP metabolism and showcase the importance of functional overlapping in bacterial signaling. Moreover, our design overcomes the native redundancy in c-di-GMP synthesis, providing a framework to dissect individual DGC functions and paving the way for targeted strategies to address bacterial adaptation and pathogenesis.

## 1. INTRODUCTION

Bacterial survival depends on the precise coordination of extracellular and intracellular signals to adapt to environmental changes [1]. These signals are conveyed through diverse small molecules that interact with complex regulatory networks, ultimately controlling bacterial growth and behavior. Cyclic dinucleotides, especially *bis*-(3′→5′)-cyclic dimeric guanosine monophosphate (cyclic di-GMP, henceforth referred to as c-di-GMP), are versatile second messengers [2], regulating numerous physiological processes in bacteria [3]. c-di-GMP, one of the earliest discovered and most widespread second messengers in bacteria, is synthesized by diguanylate cyclases (DGCs) with GGDEF domains and degraded by phosphodiesterases (PDEs) containing EAL or HD-GYP domains. However, not all GGDEF-containing proteins (GCPs) are active DGCs; some lack enzymatic activity and instead regulate c-di-GMP signaling pathways [4]. Similarly, certain EAL- or HD-GYP-containing proteins do not function as PDEs but fulfill other roles [5]. DGCs and PDEs respond to internal and external stimuli, adjusting c-di-GMP levels and initiating specific regulatory responses through interactions with effector molecules [6–8].

A key cellular process regulated by c-di-GMP is the reversible switch between motile and sessile states as bacteria transition from planktonic forms to biofilms [9]. Biofilms are surface-attached bacterial communities embedded in a self-produced matrix [10–14]. Beyond controlling the motile-sessile transition, c-di-GMP influences essential bacterial processes, including virulence, development, morphogenesis, and stress responses [15]. Given the central role of c-di-GMP, DGCs and PDEs are the subject of intense research and constitute promising targets for the design of novel drugs to combat antimicrobial resistance [16].

Bacterial species display a variable number of c-di-GMP-metabolizing enzymes. Obligate pathogens often have fewer of these enzymes compared to free-living species [15,17]. The apparent redundancy of such proteins in free-living bacteria supports the regulatory flexibility needed to adapt to fluctuating environments. The opportunistic pathogen *Pseudomonas aeruginosa*, for instance, encodes up to 40 GGDEF-, EAL-, or HD-GYP-domain proteins, or dual-domain variants, many of which are known to be involved in c-di-GMP metabolism. *P*. *aeruginosa* causes severe infections in individuals with compromised defenses, burns, or cystic fibrosis, often leading to high morbidity and mortality [18,19]. Treating *P. aeruginosa* infections is especially challenging due to a wide array of virulence factors and a remarkable capacity to resist antibiotics [20–23]. Furthermore, *P*. *aeruginosa* forms robust biofilms that evade immune clearance and impair antibiotic efficacy. The complexity of the c-di-GMP signaling network, which regulates exopolysaccharide production, biofilm formation, motility, virulence, and antibiotic resistance, underlies the adaptability and pathogenicity of this species [6,3,24].

Several approaches have been used to explore the role of c-di-GMP metabolizing enzymes in *P*. *aeruginosa*, including targeted deletions to create single mutants through specific or random (transposon) mutagenesis [25–28]. However, these methods failed to address redundancy or compensatory effects among enzymes. For example, single deletions of these proteins often produce no detectable phenotypic changes. Complementary strategies that examine individual DGCs in the absence of other c-di-GMP-synthesizing and processing enzymes would provide valuable insights into their specific roles and interactions by establishing interference-free systems. This purpose has been fulfilled by mutating genes that encode c-di-GMP-synthesizing proteins in *Salmonella enterica* [29], *Caulobacter crescentus* [30], *Sinorhizobium* (*Ensifer*) *meliloti* [31], and *Dickeya zeae* [32], with the removal of 12, 11, 15, and 16 GCPs, respectively. However, depleting all GCPs in the complex *P*. *aeruginosa* c-di-GMP signaling network remains a significant challenge, requiring extensive genome engineering that traditional methods cannot achieve efficiently. CRISPR-associated technologies offer new possibilities in bacterial genome engineering [33–35], enabling multiple genetic modifications in a cost- and time-efficient manner.

In this study, we applied a CRISPR/Cas9-based multiplex genome-editing tool to target all 32 GCP-encoding genes in *P. aeruginosa* UCBPP-PA14. We then examined how the absence of these proteins affected multiple physiological and metabolic traits, including cell growth, biofilm formation, virulence factor expression, competitiveness, antibiotic resistance, and motility. Genetic complementation with selected DGCs in the resulting mutant, termed *P*. *aeruginosa* PA14Δ32, revealed diverse effects on c-di-GMP-related traits. These results shed new light on the multifaceted role of GCPs under conditions where all such proteins are disrupted, overcoming pathway redundancy and emphasizing the complexity and adaptability of c-di-GMP metabolism in this model opportunistic pathogen.

## 2. EXPERIMENTAL PROCEDURES

### 2.1. Molecular biology techniques and construction of *P*. *aeruginosa* PA14Δ32

The bacterial strains and oligonucleotide sequences used in this study are listed in **Tables S1** and **S2** in the Supporting Information, respectively. All kits and enzymes were used according to the manufacturer’s recommendations. Plasmid and DNA purifications were performed using the NucleoSpin Plasmid EasyPure mini kit and NucleoSpin Gel and PCR clean-up mini kit (Macherey-Nagel, Düren, Germany). PCR amplifications employed Phusion *U* high-fidelity DNA polymerases (Thermo Fisher Scientific, Waltham, MA, USA), and One*Taq* Quick-Load 2× master mix (New England Biolabs, Ipswich, MA, USA). All restriction enzymes were FastDigest (Thermo Fisher Scientific), while the USER enzyme (New England Biolabs) was used for USER-cloning procedures. Sanger sequencing was carried out using the Mix2Seq kit (Eurofins Genomics GmbH, Ebersberg, Germany).

The construction of the pBEC/pMBEC plasmid for base-editing and genetic modifications of *P*. *aeruginosa* followed the methodology of Volke et al. [36]. Spacer sequences (20-nt) were designed using the CRISPy web service [37] based on the GenBank sequence (NC_008463.1) of *P*. *aeruginosa* UCBPP-PA14 [38–41], henceforth referred to as *P*. *aeruginosa* PA14 (**Table S1**). Spacers used in the study are listed in **Table S2**. The synthetic spacers were cloned into pBEC/pMBEC vectors *via* Golden Gate assembly [42], and each construct was verified through Sanger sequencing. For genome editing, *P*. *aeruginosa* PA14 was electroporated with pBEC/pMBEC plasmids containing spacers targeting genes encoding diguanylate cyclases. Cells were incubated in lysogeny broth (LB) for 3 h to recover, followed by inoculation of 100 μL into 10 mL of LB with 30 μg mL^−1^ gentamicin and incubation overnight (ON, ca. 16 h) at 37°C with agitation at 200 rpm. To cure the base-editing plasmid, 100 μL of the culture was transferred to 10 mL of LB containing 10% (w/v) sucrose and incubated ON. Dilutions of the culture were plated on LB agar to isolate individual colonies. For each base-editing event, the genomes of 5–10 randomly selected colonies were sequenced to confirm the introduction of premature *STOP* codons in the target genes.

### 2.2. Bacterial strains, plasmids, and culture conditions

*Escherichia coli* and *P. aeruginosa* were routinely incubated at 37°C in LB. Routine cloning was performed in laboratory *E*. *coli* strains [43–45]. Whenever necessary, antibiotics were used at the following concentrations: 10 μg mL^−1^ gentamicin, 100 μg mL^−1^ ampicillin, 50 μg mL^−1^ kanamycin, 30 μg mL^−1^ chloramphenicol, and 50 μg mL^−1^ streptomycin for *E. coli*; and 30 μg mL^−1^ gentamicin, 25 μg mL^−1^ tetracycline, and 250 μg mL^−1^ streptomycin for *P*. *aeruginosa*. Bacterial growth was monitored by measuring the optical density at 600 nm (OD_600_); growth parameters were calculated as described by Wirth et al. [46].

### 2.3. Whole-genome sequencing and identification of off-target mutations

Whole-genome sequencing was conducted using an Illumina Novaseq 6000 platform (Novogene Co., San Jose, CA, USA). Reads were mapped against the *P. aeruginosa* PA14 reference genome using the Geneious platform (Dotmatics, Boston, MA, USA). Sequences were trimmed and normalized before alignment. Single nucleotide polymorphisms (SNPs) and small insertions or deletions (indels) were identified *via* variant calling with a variant frequency threshold of 0.4 for all samples. A lower threshold of 0.1 was applied to the negative control to detect pre-existing nucleotide polymorphisms in the parental strain [47].

### 2.4. Phenotypic characterization

#### 2.4.1. Colony morphology

Strains were cultivated ON at 37°C with agitation in LB. Subsequently, cultures were diluted and plated on LB agar to yield 30-50 colonies per plate, followed by incubation for 16 h at 37°C. Antibiotics or inducers were added to LB and LB agar plates as required for the specific assay. Colony morphology pictures were then taken as described elsewhere [48].

#### 2.4.2. Motility assays

Swarming motility assays were conducted on M8 plates [49] containing 0.5% (w/v) agar and incubated for 48 h at 37°C, following the protocol by Caiazza et al. [50]. Swimming motility assays were performed in tryptone plates with 0.3% (w/v) agar, incubated for 24 h at 37°C. Twitching motility assays were carried out in LB agar plates containing 1% (w/v) agar, incubated for 24 h at 37°C and then for 48 h at room temperature, as described by Smania et al. [51].

#### 2.4.3. Biofilm formation

Biofilm formation was quantified using the crystal violet staining (CVS) method in a 96-well microtiter plate [52]. Replicates of the different strains were incubated ON, and cultures were adjusted to a cell density of 1.5×10^6^ UFC mL^−1^. A 100-μL aliquot od each culture was seeded in triplicate into a 96-well microtiter plate and incubated at 37°C for 16 h. After incubation, biofilms were stained with crystal violet, and the absorbance was measured at 595 nm in an Epoch^TM^ Microplate Spectrophotometer (BioTek Instruments Inc., Winooski, VT, USA).

Biofilm imaging of *P. aeruginosa* was performed as described by Moyano et al. [53]. Bacterial strains were grown in 8-well Lab-Tek chamber coverglass systems (Nunc^TM^, Thermo Fisher Scientific). Chambers were inoculated with 5×10^5^ cells in 250 μL of fresh medium and incubated for 24 h at 37°C. Biofilms were stained using the LIVE/DEAD BacLight^TM^ bacterial viability kit (Molecular Probes Inc., Eugene, OR, USA), applying the SYTO 9 green-fluorescent nucleic acid stain according to the manufacturer’s protocol. Images were captured with an FV1200 confocal microscope (Olympus, Tokyo, Japan) using 60× oil super-corrected UPLXAPO60XO objective lens. Cell detection was performed with the default configuration of FV10 software (ASW 4.0; Evident, Waltham, MA, USA) for green emission fluorophores (SYTOX^TM^ green nucleic acid stain; Sigma-Aldrich Co., St, Louis, MO, USA).

#### 2.4.4. Virulence assays

Rhamnolipid production was assessed by observing red blood cell lysis on sheep blood agar plates (Britania S.A., Buenos Aires, Argentina). Plates were inoculated with 5 μL of bacterial culture and incubated at 37°C for 96 h. Extracellular protease activity was evaluated using milk agar plates containing 1% (w/v) agar. Bacteria were first grown aerobically in tryptic soy broth at 37°C, after which 5 μL of culture was spotted on milk agar plates and incubated overnight at 37°C. Degradation halos and colony diameters were measured and quantified using ImageJ software (Fiji; [54]. Pyoverdine production was assessed in King’s B medium [51] after cultures were incubated at 37°C for 24 h with agitation at 220 rpm. A 1-mL aliquot of the culture was centrifuged at 10,000×*g* for 10 min, and the absorbance of the supernatant was measured at 360 nm. Pyocyanin production was measured in LB. Cultures were grown for 48 h at 220 rpm, followed by centrifugation of 2 mL at 10,000×*g* for 10 min. Pyocyanin was extracted from the supernatant with chloroform/HCl, and the absorbance of the extract was measured at 595 nm.

#### 2.4.5. Nematode paralysis assay

Nematode paralysis assays were conducted following the method by Luján et al. [55]. In brief, 10 mL aliquots of 1/10 dilutions of ON brain heart infusion (BHI) cultures were spread onto 5.5-cm diameter plates containing 10 mL of BHI with 1.2% (w/v) agar. Plates were incubated for 24 h at 30°C. *Caenorhabditis elegans* var. Bristol strain N2 nematodes were collected from stock plates using M9 buffer (containing 20 mM KH_2_PO_4_, 40 mM Na_2_HPO_4_, 90 mM NaCl, and 1 mM MgSO_4_), and a 50-μL aliquot containing ca. 20-40 adult worms was spotted onto the *P. aeruginosa* lawn. Plates were incubated for 6 h at room temperature with lids on, after which the number of live nematodes was recorded. Worms were deemed paralyzed if they failed to move spontaneously and showed no response to mechanical stimulation. *E. coli* strain OP50 [56] was included as non-virulent control, with each assay performed at least in triplicate.

#### 2.4.6. Transmission electron microscopy

For transmission electron microscopy (TEM), strains were grown ON at 37°C on LB agar plates. Cells were then suspended in 1× PBS (containing 137 mM NaCl, 2.7 mM KCl, 10 mM Na_2_HPO_4_, and 1.8 mM KH_2_PO_4_, pH = 7.4) by scraping the biomass from the agar plate surface. TEM imaging was carried out at the Transmission Electron Microscopy Services of Instituto de Investigaciones en Ciencias de la Salud (INICSA)-CONICET, Centro de Microscopía Electrónica (Facultad de Ciencias Médicas, Universidad Nacional de Córdoba, Córdoba, Argentina). Formvar-coated nickel grids were floated on a mixture of bacteria suspension and a 2% (v/v) H_3_PW_12_O_40_ solution in a 1:1 ratio for 1 min. The grids were then rinsed briefly with distilled water, and excess liquid was removed using the edge of filter paper. After air drying for 5 min, the specimens were examined in a Zeiss LEO906 TEM (Carl Zeiss AG, Oberkochen, Germany) operated at an accelerating voltage of 80 kV and photographed with a Megaview G3 camera (EMSIS GmbH, Münster, Germany).

#### 2.4.7. Expression of DGC genes for complementation assays

The pJN105 vector [57] carrying wild-type copies of *PA14_16500* or *PA14_23130* (plasmids pJN_*wspR* and pJN_*23130*, respectively; **Table S1**) was used for the expression of the DGC-encoding genes in complementation assays. In these constructs, the expression of *PA14_16500* and *PA14_23130* was driven by an arabinose-inducible expression system [58,59]. For routine complementation assays, transformed cells were cultured in LB or on LB agar plates supplemented with 0.1-1% (w/v) arabinose. Additionally, a wild-type copy of *fimX* (*PA14_65540)* was cloned into the pMBLe vector, generating plasmid pMBLe_*fimX* [20], with *fimX* expression induced with IPTG at 1 mM.

#### 2.4.8. Competence with S*taphylococcus aureus*

Bacteria were cultured aerobically in tryptone soy agar 37°C. A saturated culture of *S. aureus* USA300 [60] was adjusted to an OD_600_ = 1, and 100 μL was spread onto a tryptic soy agar plate. A 5-μL suspension of *P. aeruginosa* PA14 or PA14Δ32 was spotted onto these plates, which were incubated ON at 37°C. Inhibition halos, along with colony diameters, were measured using the ImageJ software (Fiji).

#### 2.4.9. Antibiotic susceptibility testing

Minimal inhibitory concentrations (MICs) were determined by using Sensititre^TM^ plates (Thermo Fisher Scientific). *P. aeruginosa* PA14 and PA14Δ32 were cultured ON in LB at 37°C under aerobic conditions. These cultures were then used in antibiotic test panels by following the manufacturer’s instructions. To do this, the OD_600_ was diluted to a 0.5 McFarland scale, and 5 μL of the suspension was transferred to commercial Mueller-Hinton broth adjusted for cation concentration. A 50-μL aliquot of this suspension was then added to each well of the Sensititre^TM^ plate. The inoculated plates were incubated ON at 37°C, and bacterial growth was determined in each well.

#### 2.4.10. Bacterial growth profile on different carbon sources

Single colonies of the strains under study were used to inoculate precultures, grown in the same medium and conditions used for subsequent growth monitoring. Precultures were incubated for 24 h. For growth monitoring, 200 μL of medium was inoculated with the corresponding preculture to an OD_600_ = 0.02 and dispensed into a 96-well plate. The cultures were incubated at 37°C in a Synergy HI plate reader (BioTek Instruments Inc.), and OD_600_ was recorded every 15 min. Media tested included M9 minimal medium supplemented with 10 mM citrate, glucose, gluconate, glycerol, succinate, or succinate [61], as well as LB.

#### 2.4.11. Bacterial tolerance to stress conditions

Single colonies of the specified strains were used to inoculate precultures, which were grown for 24 h. Growth was monitored in 1 mL of the corresponding medium, inoculated with the preculture to an OD_600_ = 0.02 and dispensed into a 96-well plate. Cultures were incubated at 37°C in a Synergy HI plate reader (BioTek Instruments Inc.), with OD_600_ measured every 15 min. pH tolerance was evaluated using LB adjusted to a pH range of 6 to 10, with phosphate buffer for pH = 6 to 9 and glycine phosphate buffer for pH = 10. Saline stress was assessed using LB containing NaCl concentrations up to 1 M. Oxidative stress analysis involved aerobic cultures in tryptic soy agar at 37°C, followed by spreading 100 μL of culture onto LB agar plates. Filter disks embedded with 10 μL of H_2_O_2_ were placed on the plates, and after ON incubation at 37°C, inhibition halos and colony diameters were measured using the ImageJ software (Fiji).

### 2.5. Determination of c-di-GMP levels through a biosensor and analytical quantification by LC-MS/MS

A fluorescent biosensor, comprising the monomeric green fluorescent protein (msfGFP) gene regulated by the *P_pelA_* promoter [62], was integrated as a Tn*7* module into the *P. aeruginosa* chromosome by tetraparental mating [63]. For this, *E. coli* PIR2 (Thermo Fisher Scientific) carrying plasmid pBG·PelA or pBG·PelA·Sm (donor strain), *E. coli* DH5α λ*pir* carrying plasmid pTnS1 (helper strain expressing the transposase), *E. coli* HB101 carrying plasmid pRK600 (helper strain), and *P. aeruginosa* (recipient strain) were mixed in LB, streaked on LB agar plates, and incubated at 37°C for 24 h [64,65]. *P. aeruginosa* clones carrying *att*Tn*7*::[*P_pelA_*(BCD2)→*msfGFP*] chromosomal integrations were selected on cetrimide agar plates containing gentamicin and confirmed by colony PCR [66]. These clones were stored as glycerol stocks, streaked on LB agar plates, and four colonies were grown in M9 minimal medium for 16 h at 37°C under the conditions specified in the text. After washing with 1× PBS and standardizing to and OD_600_ = 1, fluorescence was measured using a Synergy HT^TM^ microtiter plate reader (BioTek Instruments Inc.).

ON cultures of both PA14 and PA14Δ32 strains were prepared in 10 mL of LB at 37°C with shaking at 200 rpm. Antibiotics were supplemented as needed to maintain plasmids during the experiments. The following day, 20 mL of LB were inoculated to an OD_600_ = 0.05 with the ON cultures and placed in a 100-mL Erlenmeyer flask. When the OD_600_ reached 0.5, the cultures were centrifuged at 4,000×*g* for 5 min at 4°C. The supernatant was discarded, and the cell pellet was quenched with 2 mL of a solution containing 40% (v/v) acetonitrile and 40% (v/v) methanol in water, acidified with 0.1 M formic acid. After vigorous vortexing, cell debris was removed by centrifugation at 13,000×*g* for 2 min. The supernatant was transferred to a fresh tube and concentrated to 0.1 mL at 30°C for 2-3 h using an Eppendorf^TM^ Concentrator Plus (Sigma-Aldrich Co.). The analysis of c-di-GMP was performed *via* LC-MS/MS as previously described [67]. Detection parameters, optimized for c-di-GMP quantification, are provided in **Table S3**.

### 2.6. Transcriptomic analysis

Strains were cultivated according to the protocol for c-di-GMP quantification. Once cultures reached an OD_600_ = 0.5, cells from 2 mL of culture were harvested by centrifugation at 4,000×*g* for 5 min at 4°C. The resulting cell pellets were flash-frozen in liquid nitrogen and stored at –80°C for subsequent analysis. RNA extraction was performed as described by Turlin et al. [68], and RNA quality was evaluated using an Agilent 2100 Bioanalyzer (RNA 6000 Nano kit; Agilent Technologies, Waldbronn, Germany), assessing RNA concentration, RNA integrity number, 28S/18S ratio, and fragment length distribution. Library construction, sequence filtering, mapping, and genome expression analysis were outsourced to BGI Genomics Co. (Hong Kong). Protein-protein interaction network and enrichment analyses of differentially expressed genes were conducted using the *STRING* web platform (https://string-db.org/; [69].

### 2.7. Data and statistical analysis

All the experiments were independently repeated at least twice, as indicated in the corresponding figure or table legend, and the mean value of the corresponding parameter ± standard deviation is presented unless otherwise indicated. When relevant, the level of significance of the differences when comparing results was evaluated by ANOVA and Tukey multiple comparison test or the Student’s *t-*test with α = 0.05.

## 3. RESULTS

### 3.1. Construction of a *P. aeruginosa* strain with disrupted GCPs

*P. aeruginosa* possesses one of the most complex c-di-GMP signaling networks among bacteria, with 40 proteins containing DGC or PDE domains. Our primary objective was to generate a *P*. *aeruginosa* strain in which all c-di-GMP–metabolizing proteins were inactivated. To achieve this goal, we adopted a multiplex CRISPR/Cas base-editing system previously developed for *Pseudomonas* species in our laboratory [36], which enables the simultaneous inactivation of multiple genes (**Fig. 1A**). This system expresses multiple gRNAs under a single promoter, which are post-transcriptionally processed by the endoribonuclease Cas6f into mature gRNAs with distinct spacers (**Fig. 1B**). A CRISPR/Cas complex fused to a cytidine base editor (CBE) catalyzes cytidine-to-thymine conversions that are permanently incorporated into the genome during DNA replication. We targeted 32 proteins predicted to contain a DGC domain (GGDEF or similar) in *P*. *aeruginosa* PA14 (**Fig. 2** and **Table S4**). Spacers were designed to target locations within the corresponding open reading frames (ORFs) to disrupt GCP genes, following established criteria [36]. These criteria required the editable cytidine to be positioned between the second and tenth positions within the editing window, while avoiding guanosine residues immediately preceding the editable cytidine (**Table S5**). By applying these criteria, all GCP-encoding ORFs distributed across the *P*. *aeruginosa* chromosome were successfully edited, achieving 100% base-editing efficiency.

**Fig. 1.**
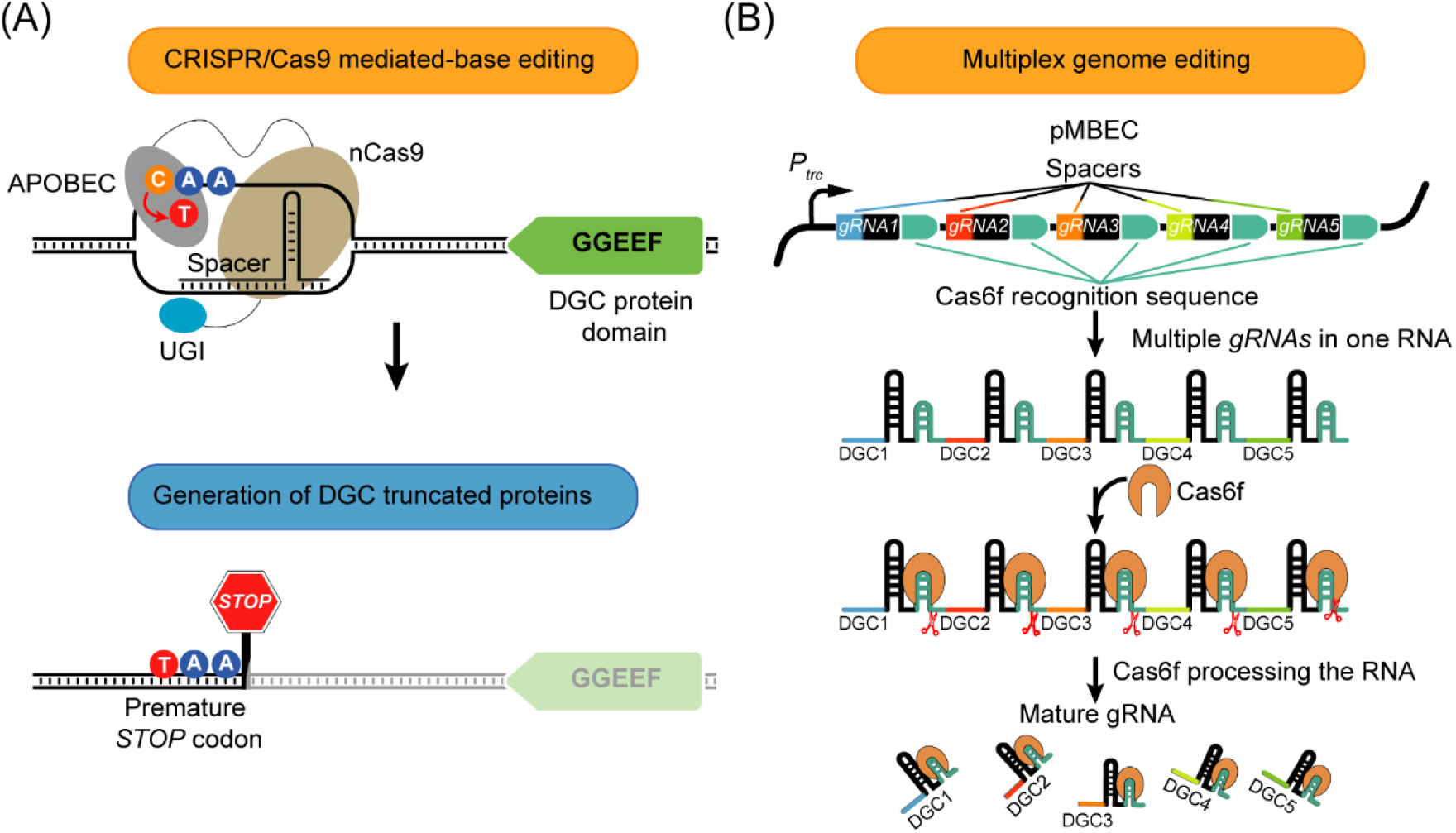
Construction of a *P. aeruginosa* strain with disrupted GCPs. **(A)** Cytidine base editor (CBE) structure. A nicking Cas9 (nCas9) guided by a single guide RNA (sgRNA) targets the cognate protospacer sequence. Upon hybridization between the sgRNA spacer and the target DNA, the complementary strand is exposed. A cytidine deaminase (APOBEC) fused to nCas9 deaminates cytidine (C) to uracil within an editing window, and during DNA replication, uridine is converted to thymidine (T). The uracil DNA glycosylase inhibitor (UGI) prevents cytidine restoration by endogenous DNA repair pathways. Spacers are designed to induce a premature *STOP* codon through the C-to-T exchange. *PAM*, protospacer adjacent motif. **(B)** RNA cassette processing by Cas6f. Each guide RNA (gRNA), cloned in pBEC/pMBEC vectors, includes a 3’ recognition site for the endoribonuclease Cas6f, which cleaves the RNA and remains bound to each gRNA post-cleavage.

**Fig. 2.**
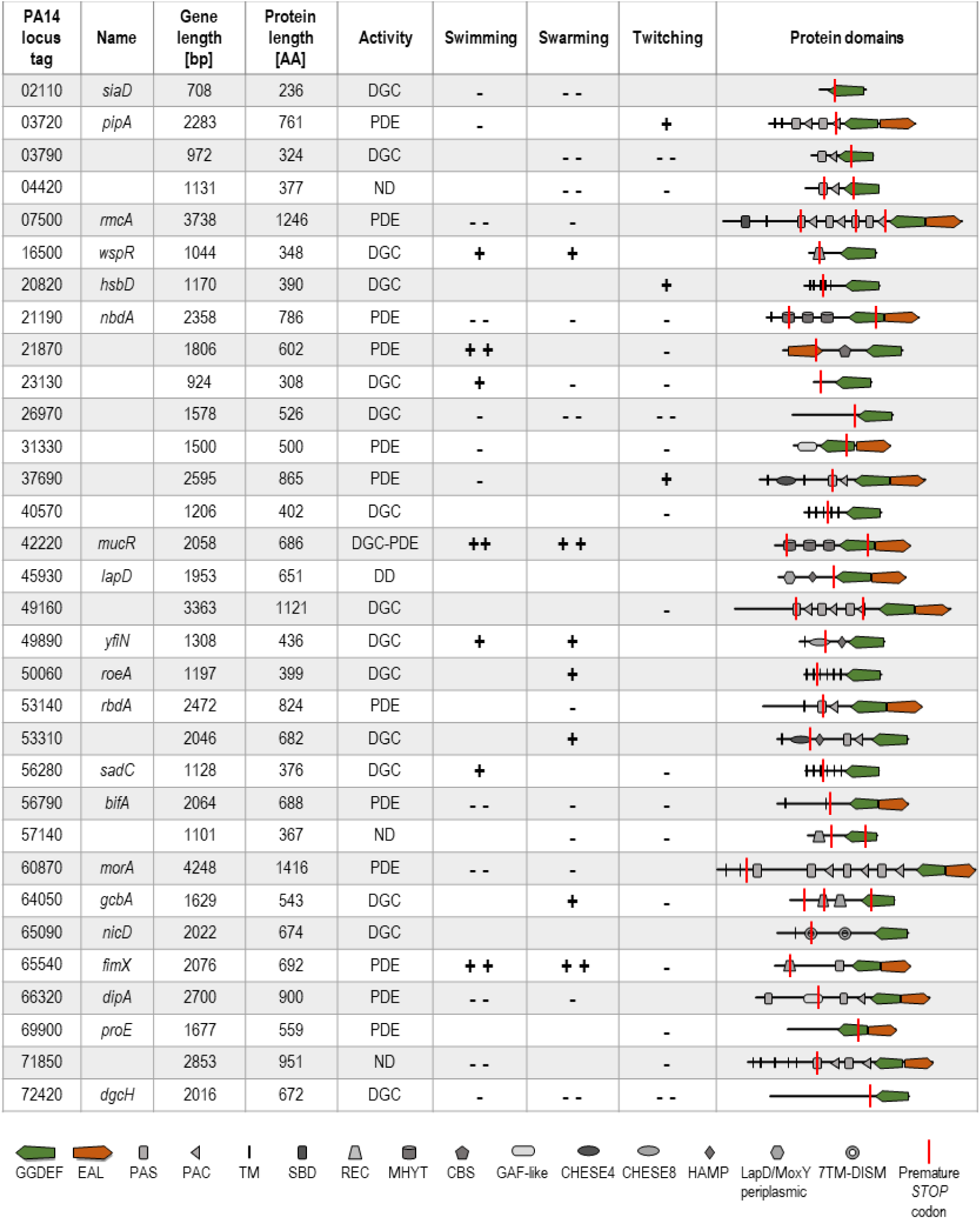
GCPs in *Pseudomonas aeruginosa* PA14. The GGDEF-containing protein (GCP) architecture and representation of premature *STOP* codons generated by CRISPR/nCas9 multiplex cytidine base-editing are shown for each GCP in *P. aeruginosa* PA14. Gene structures are not drawn to scale. Data on DGC activity, swimming, swarming, and twitching data obtained from Ha and O’Toole [75] and Kulasakara et al. [27]; see also **Table S4**. Symbols (+, ++, –, – –) reflect the degree of motility changes observed in single mutants analyzed by Ha et al. [25]. Empty boxes indicate no discernible motility effect. *Abbreviations*: *AA*, amino acids; *GGDEF*, diguanylate cyclase domain; *EAL*, phosphodiesterase domain; *PAS*, signal transduction domain; *TM*, inner membrane transmembrane domain; *SBD*, periplasmic substrate-binding domain; *REC*, response regulator domain; *MHYT*, integral membrane protein domain; *CBS*, cystathionine β-synthase domain; *GAF-like*, c-di-GMP–specific phosphodiesterases, adenylyl cyclases, and FhlA domain; *CHESE4* and *CHESE8*, periplasmic sensor domains; *HAMP*, histidine kinases, adenylyl cyclases, methyl binding proteins, phosphatases; *LapD/MoxY periplasmic*, N-terminal periplasmic domain of the LapD and MoxY receptor proteins; *7TM-DISM*, signal transduction domain with seven transmembrane receptors with diverse intracellular signaling modules.

*P. aeruginosa* DGC proteins typically have a GGDEF domain at the C-terminus, while in most dual GGDEF/EAL-domain proteins, the GGDEF domain is followed by the EAL domain near the C-terminus (**Fig. 2**). This arrangement enabled the introduction of premature *STOP* codons in the ORF of each gene upstream of the GGDEF motif (**Fig. 2**). The ORFs *PA14_31330*, and *PA14_69900*, predicted to encode dual EAL-GGDEF proteins previously identified as phosphodiesterases rather than diguanylate cyclases [70,71], were modified by partially interrupting the GGDEF domain with *STOP* codons. For *PA14_21870*, the only protein where the EAL domain is N-terminal to the GGDEF motif, the target cytidine residue was situated between the two domains. In a few other cases (*PA14_04420*, *PA14_07500*, and *PA14_64050*), the ORFs contained more than one targeted cytidine residue.

Derivatives of the base-editing pBEC/pMBEC vectors [36] carrying the required gRNAs to modify all 32 targets were assembled (**Table S1**) and introduced into *P*. *aeruginosa* PA14 through iterative editing cycles (**Fig. S1A**). After each cycle, colony PCR was performed on randomly selected isolates, and amplicon sequencing confirmed successful edits, enabling the selection of partial mutants for subsequent rounds of base-editing. When a given spacer proved inefficient at editing its target gene, alternative spacer designs were tested and implemented (**Table S2**). By the end of this iterative base-editing process, whole-genome sequencing verified the successful editing of all 32 targets. This genome-engineering effort produced *P*. *aeruginosa* PA14Δ32, a mutant strain in which all known c-di-GMP–synthesizing enzymes (i.e., *wspR*, *yfiN*, *morA*, PA14_72420, *rbdA*, *bifA*, PA14_71850, PA14_50060, PA14_53310, PA14_65090, PA14_20820, PA14_26970, PA14_40570, PA14_21870, PA14_31330, PA14_37690, PA14_45930, PA14_65540, PA14_03720, PA14_69900, PA14_04420, PA14_23130, PA14_02110, PA14_03790, PA14_42220, PA14_49160, PA14_66320, PA14_21190, PA14_07500, PA14_57140, PA14_56280, and PA14_64050; **Table S1** and **Fig. S1B**) were disrupted.

### 3.2. *P*. *aeruginosa* PA14Δ32 exhibits stable physiology across growth conditions despite substantially decreased c-di-GMP levels

The PA14Δ32 mutant displayed a plateau-like colony morphology with plaque-like clearing zones at the centers, contrasting with the regular colony phenotype of the wild-type strain on LB agar (**Fig. 3A**). Growth rates in complex and minimal media supplemented with various carbon and energy sources were lower than those of the parental strain under all tested conditions (**Fig. 3B**). However, the extent of this reduction suggests a relatively limited disruption to the overall bacterial physiology due to the lack of DGC proteins. The most pronounced growth rate decrease (ca. 35%) was observed in cultures containing gluconeogenic substrates, while cultures in LB exhibited only a slight reduction (ca. 14%) compared to the wild-type strain. Furthermore, strain PA14Δ32 was more sensitive to oxidative stress, as evidenced by larger inhibition halos following H_2_O_2_ exposure (**Fig. 3C**). No significant differences were observed in responses to variations in pH (**Fig. S2A**) or osmolarity (**Fig. S2B**), tested in the presence of NaCl [72].

**Fig. 3.**
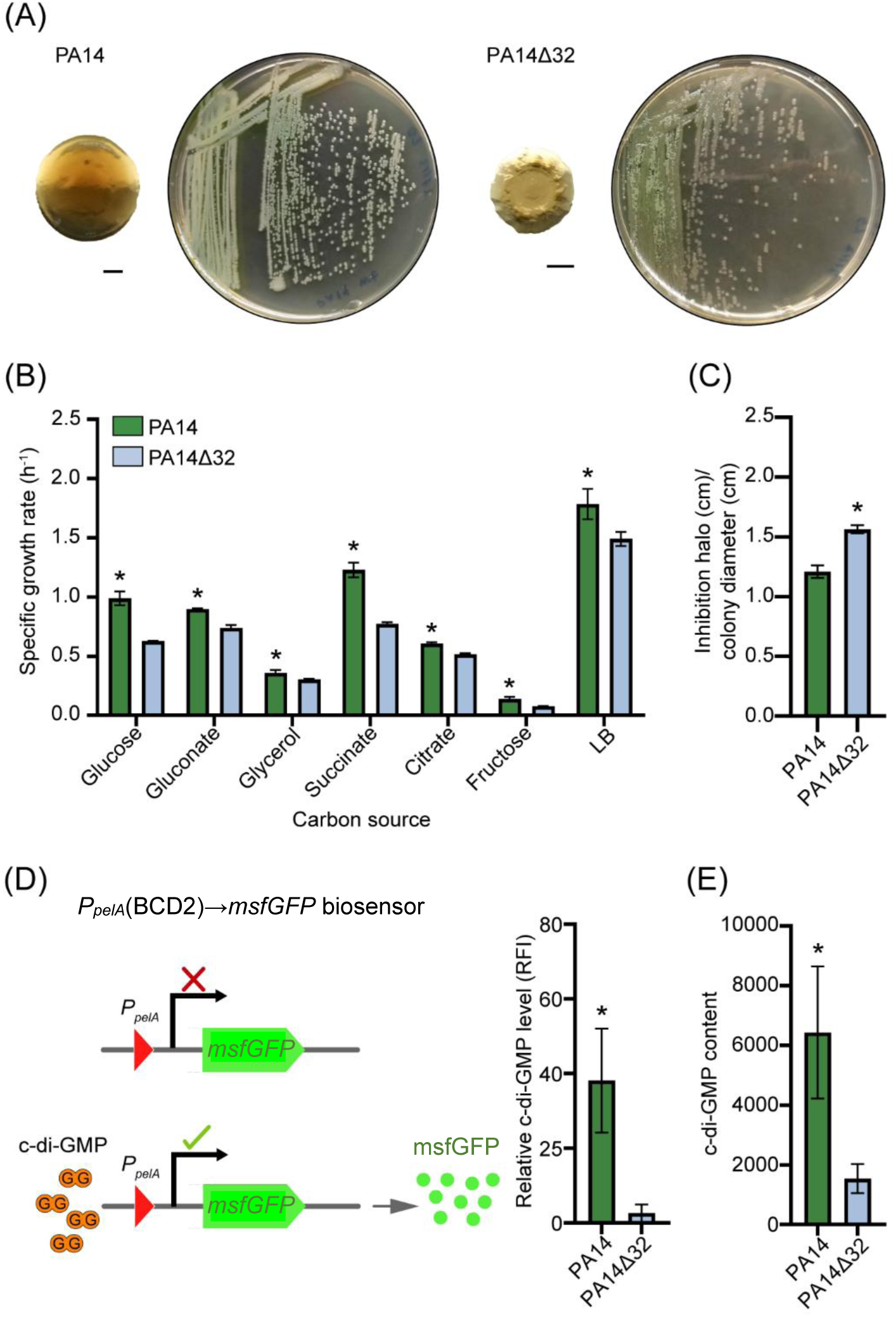
Strain PA14Δ32 maintains consistent physiology across growth conditions despite substantially reduced c-di-GMP levels. **(A)** Colony morphology and streak culture images of PA14 and PA14Δ32 strains (scale bars = 2 mm). LB agar plates were incubated for 24 h at 37°C, revealing autolysis in PA14Δ32. **(B)** Specific growth rate comparison between PA14 and PA14Δ32 strains on various carbon sources. Cultures were inoculated in minimal medium supplemented with different carbon sources and grown for 24 h at 37°C with agitation. **(C)** Growth under oxidative stress conditions. Bacteria were cultured in tryptic soy medium, spread on LB agar plates, and exposed to H_2_O_2_ in filter-paper disks embedded with 10 μL of H_2_O_2_. Inhibition halos and colony diameter were measured after overnight incubation at 37°C. Results represent average values ± standard deviation from three independent experiments. **(D)** c-di-GMP levels and architecture of the biosensor. The monomeric and superfolder green fluorescent protein (msfGFP) is under transcriptional control of the c-di-GMP–responsive *P_pelA_* promoter from the *P*. *aeruginosa pel* operon. Strains were cultured in minimal medium supplemented with glucose and acid casein hydrolysate for 24 h and standardized to the same optical density at 600 nm (OD_600_). Results represent the mean relative fluorescence intensity (RFI) ± standard deviations from triplicate measurements from least two independent experiments. **(E)** c-di-GMP levels quantified *via* LC-MS/MS. Cultures of both strains were grown in LB and adjusted to the same OD_600_. Results represent the average c-di-GMP content ± standard deviations relative to wild-type *P*. *aeruginosa* PA14, based on three independent experiments. In all cases, statistical significance with *p* < 0.05 is indicated by an asterisk (*) symbol (Student’s *t*-test).

Intracellular c-di-GMP levels were semi-quantitatively assessed using a c-di-GMP–responsive fluorescent biosensor [63]. This system places the gene encoding msfGFP under the transcriptional control of the c-di-GMP–responsive *P_pelA_* promoter from *P*. *aeruginosa* (**Fig. 3D**). In its native context, this promoter regulates the expression of the *pel* operon, involved in exopolysaccharide biosynthesis [73]. The biosensor-based measurements revealed negligible residual c-di-GMP levels in the PA14Δ32 strain (**Fig. 3D**). In contrast, LC-MS/MS c-di-GMP quantification revealed significantly reduced (>80%) dinucleotide concentrations in cell extracts of *P*. *aeruginosa* PA14Δ32 (**Fig. 3E**), probably due to the lower detection limit as compared to fluorescence-based biosensors. This result suggests that some targeted enzymes may retain partial activity or that cryptic pathways contribute to c-di-GMP synthesis in the PA14Δ32 background. Based on these observations, we focused on characterizing this *P*. *aeruginosa* mutant with markedly reduced c-di-GMP levels.

### 3.3. Differentially expressed genes upon GCPs disruption in strain PA14Δ32

To examine the impact of GCP disruption on gene expression, we performed a genome-wide transcriptomic analysis using deep RNA sequencing on exponentially growing cultures of strain PA14Δ32 grown in LB. Differential gene expression was identified by setting thresholds of log_2_ of the fold-change (FC) > 2 or < –2 as an indication of transcriptional up- or down-regulation, respectively. The analysis revealed significant alterations in the transcription of 225 genes in strain PA14Δ32 compared to the parental strain, representing 4% of the gene repertoire of *P. aeruginosa*. Among these, 85% (192 genes) showed down-regulation, with log_2_(FC) values < –2, while 15% (33 genes) were up-regulated. Detailed transcriptional changes are listed in **Table S6**, and a volcano plot summarizing the data is presented in **Fig. 4A**.

**Fig. 4.**
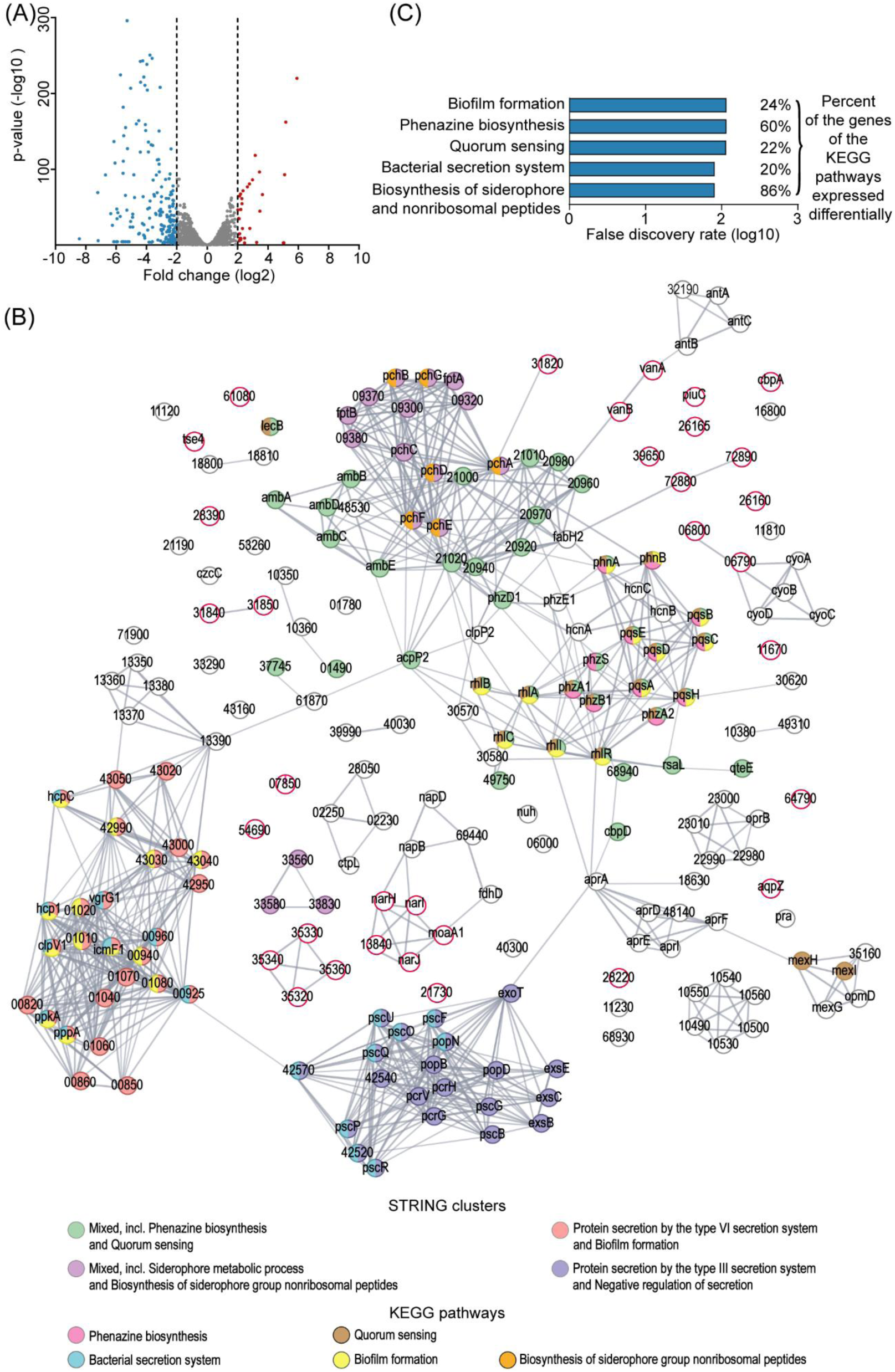
Differentially expressed genes upon GCPs disruption in strain PA14Δ32. PA14 and PA14Δ32 strains were cultured with agitation at 37°C until mid-exponential phase (OD_600_ = 0.5). Cells were harvested for RNA sequencing to identify differentially expressed genes in PA14Δ32. Genes with *p* < 0.05 and a log_2_(fold-change) ≥ 2 (upregulated) or ≤ –2 (downregulated) compared to wild-type *P*. *aeruginosa* PA14 were selected. **(A)** Volcano plot of RNA sequencing analysis, where blue circles represent downregulated genes and red circles represent upregulated genes. **(B)** Protein-protein interaction network of differentially expressed genes, constructed using the *STRING* web tool [69]. Clusters were organized based on protein interactions and their roles in specific cellular processes. Red-lined circles indicate upregulated genes, with PA14 loci labeled by their specific numbers. **(C)** Kyoto Encyclopedia of Genes and Genomes (KEGG, [74]) pathway enrichment analysis of differentially expressed genes. The percentage of genes deregulated in each KEGG pathway in PA14Δ32 is displayed on the right side of the bars. False discovery rates (FDRs) describe pathway enrichment significance, with values corrected for multiple testing using the Benjamini–Hochberg procedure, as implemented in the *STRING* web tool [69].

None of the disrupted GCPs showed significant changes in transcription, except for *PA14_21190* (**Table S7**), which displayed a modest perturbation with a log_2_(FC) = –2. To investigate the biological functions or metabolic pathways impacted by GCP disruption, we analyzed differentially expressed genes using the *STRING* database [69]. A protein-protein interaction network was constructed based on physical interactions and functional associations derived from curated sources, including Kyoto Encyclopedia of Genes and Genomes (KEGG) pathways [74]. In this network, the differentially expressed genes were organized into four primary clusters, all comprising downregulated genes (**Fig. 4B**). These clusters included (i) phenazine biosynthesis and quorum sensing, (ii) siderophore metabolism and biosynthesis, (iii) protein secretion *via* the type VI secretion system and biofilm formation, and (iv) protein secretion *via* the type III secretion system and negative regulation of secretion (**Data S1**). Together, these clusters accounted for 53% of the downregulated functions and 45% of all differentially expressed genes. Notably, they aligned with five metabolic pathways affected by the transcriptionally downregulated genes in strain PA14Δ32: biofilm formation, phenazine biosynthesis, quorum sensing, bacterial secretion systems, and siderophore and non-ribosomal peptide biosynthesis (**Fig. 4C** and **Data S1**). The most significant transcriptional changes occurred in phenazine and siderophore biosynthesis, with 60% and 86% of pathway genes, respectively, being downregulated. Analysis of off-target mutations revealed no changes in genes associated with these metabolic pathways (**Data S2**). Taken together, these findings demonstrate that disrupting GCP in *P*. *aeruginosa*, which reduces c-di-GMP levels, significantly attenuates key mechanisms governing biofilm formation and virulence. The experimental validation of the transcriptional analysis results is detailed below.

### 3.4. Biofilm formation is suppressed in strain PA14Δ32

c-di-GMP is a critical regulator of biofilm formation, with elevated intracellular levels linked to sessile cells and increased biofilm production [75]. Biofilm formation was assessed using the standard crystal violet staining (CVS) method [52] to compare the wild-type strain with the PA14Δ32 mutant. The wild-type strain had an average biofilm index of 0.5 ± 0.2, reflecting robust biofilm formation (**Fig. 5A** and **5B**). In contrast, the PA14Δ32 mutant exhibited no detectable biofilm formation under these conditions.

**Fig. 5.**
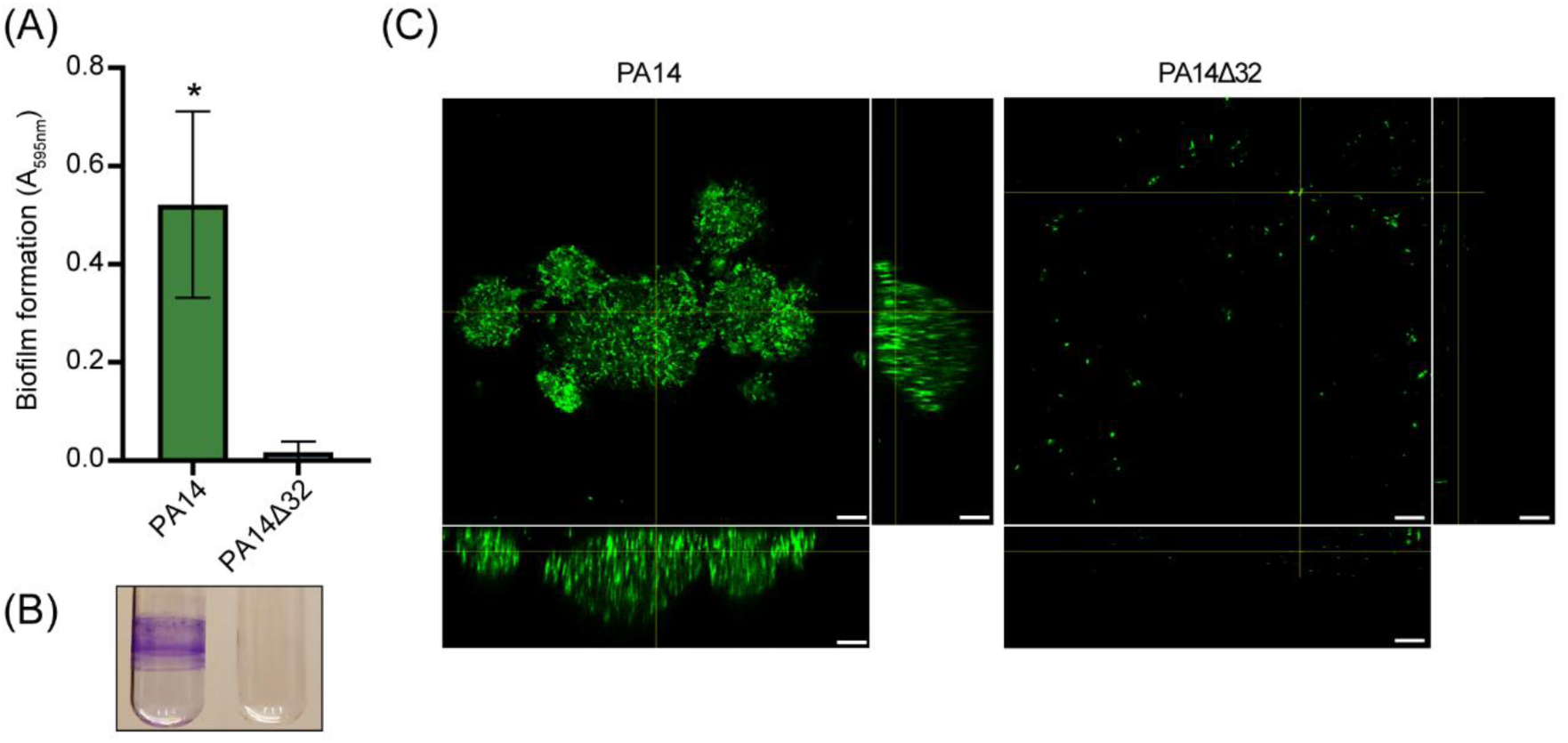
Biofilm formation is abolished in strain PA14Δ32. **(A)** Biofilm production in PA14 and PA14Δ32 strains was quantified after 24 h of growth at 37°C in polystyrene microtiter plates under static conditions. Biofilm was quantified by measuring the absorbance at 595 nm (A_595 nm_) following crystal violet staining and elution with ethanol. Results represent averages ± standard deviations from quadruplicate measurements in at least three independent experiments. Statistical significance with *p* < 0.05 is indicated by an asterisk (*) symbol (Student’s *t*-test). **(B)** Images of crystal violet-stained biofilms of *P*. *aeruginosa* PA14 and PA14Δ32 after overnight growth in LB at 37°C with agitation. **(C)** Confocal laser scanning microscopy of biofilm structures in PA14 and PA14Δ32 strains after 24 h of growth in 8-well Lab-Tek chamber slides stained with SYTO 9. Images are representative of four observations from at least two independent experiments. Scale bar = 10 μm.

Confocal laser scanning microscopy showed that the PA14Δ32 mutant lacks a defined biofilm structure, while the wild-type strain formed a well-organized biofilm. Biofilm formation by strain PA14Δ32 was virtually suppressed, with only sparse surface-attached bacteria observed in the preparation (**Fig. 5C**). These findings support the conclusion that GCP proteins orchestrate the shift from planktonic to multicellular lifestyles in *P. aeruginosa* but are not essential for cell viability. Additionally, the minimal c-di-GMP levels detected in the mutant (< 20% of those in the parental strain) appear insufficient to support biofilm production. Building on these observations, we explored other phenotypic traits potentially affected by altered c-di-GMP metabolism.

### 3.5. The GCP-disrupted mutant maintains normal swimming but lacks swarming and twitching motility

*P. aeruginosa* uses two surface structures, a single polar monotrichous flagellum and polar type IV pili, to facilitate movement. These organelles enable three distinct motility types: (i) swimming in liquid environments, driven by the flagellum, (ii) twitching across solid surfaces through pilus extension and retraction, and (iii) swarming on semisolid surfaces, which depends on flagellar stators and rhamnolipid production [50]. The relationship between c-di-GMP levels and motility involves complex interactions [76]. Mutations in enzymes regulating c-di-GMP turnover lead to varying effects on motility, from enhancement to complete movement inhibition [75]. Based on this background, the next objective was examining motility in the PA14Δ32 mutant.

The PA14Δ32 mutant retained swimming motility comparable to the wild type (**Fig. 6A**), indicating that disrupting GCPs does not significantly impact flagellum assembly or function. However, *P*. *aeruginosa* PA14Δ32 exhibited no swarming or twitching motility (**Fig. 6B** and **6C**), suggesting that one or more GGDEF-domain proteins are crucial for these specific behaviors. Transmission electron microscopy (TEM) confirmed the presence of a polar flagellum in strain PA14Δ32, structurally resembling the wild type (**Fig. 6D**). Notably, transcription of the structural *rhl* genes, which encode key components involved in rhamnolipid production (RhlABC), was significantly downregulated in the mutant (**Table S6**)—accounting for the swarming deficiency. Additionally, the inactivation of *fimX* (*PA14_65540*), a dual-domain GGDEF-EAL protein critical for type IV pilus biogenesis and assembly [77], helps explaining the observed twitching failure. In this sense, the gene encoding FimX was the final base-edit during strain construction (**Fig. S1A**), allowing its function to be tested independently of the other 31 GGDEF-domain proteins. This mutant strain, termed *P*. *aeruginosa* PA14Δ31 (**Table S1**), displayed normal twitching motility and pili (**Fig. 6C** and **6D**), comparable to the wild type. Complementation of *fimX* in the PA14Δ32 background with plasmid pMBLe_*fimX* restored both twitching motility and normal piliation levels (**Fig. 6C** and **6D**)— similar to the observations in the wild-type strain and the PA14Δ31 mutant.

**Fig. 6.**
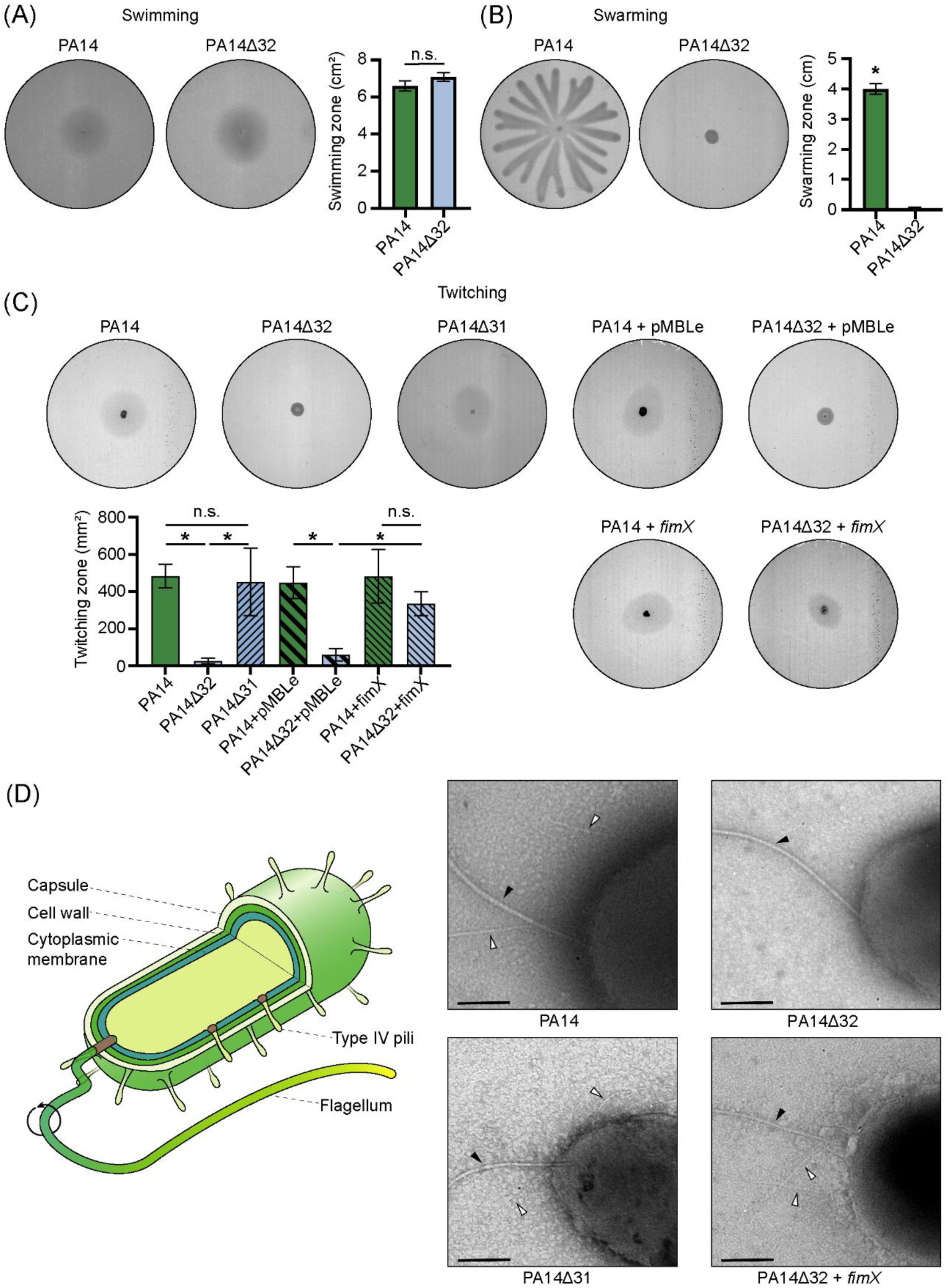
The GCP-disrupted mutant maintains normal swimming but lacks swarming and twitching motility. Analysis of **(A)** swimming and **(B)** swarming motility in *P*. *aeruginosa* PA14 and PA14Δ32. **(C)** Twitching motility analysis in strains PA14, PA14Δ32, PA14Δ31, and PA14Δ32 transformed with plasmids pMBLe (empty vector, used as a control) or pMBLe_*fimX* (+ *fimX*). Quantification in panels **(A)**, **(B)**, and **(C)** was based on the diameters of the covered areas. Measurements were performed in quadruplicate across three independent experiments and analyzed using ImageJ software. Results represent averages ± standard deviations. Statistically significant differences (*p* < 0.05) were determined using the Student’s *t*-test; the asterisk (*) symbol indicates significance for swarming, *n*.*s*. indicates not significant for swimming. Twitching motility was analyzed using one-way ANOVA followed by Tukey’s post hoc test, with the asterisk (*) symbol denoting significance at *p* < 0.05. **(D)** Schematic representation of a *P*. *aeruginosa* cell showing the flagellum and pili, accompanied by transmission electron microscopy (TEM) images used to visualize surface pili and flagella in strains PA14, PA14Δ32, PA14Δ31, and PA14Δ32 + *fimX*. White arrows indicate pili, and black arrows indicate flagella. Scale bar = 200 nm.

Taken together, these findings highlight the critical role of FimX among GGDEF-domain proteins in supporting twitching motility. Remarkably, twitching persisted even with significantly reduced c-di-GMP levels. These results prompted further experiments to assess whether the phenotypic changes in the GCP-depleted strain could impact virulence.

### 3.6. Strain PA14Δ32 has reduced production of virulence factors, while its antibiotic resistance profile remains unaffected

Virulence traits in *P. aeruginosa* are tightly regulated by different GCPs through modulation of c-di-GMP levels [27]. Transcriptomic analysis of the PA14Δ32 strain revealed a marked downregulation of key components in both the *rhl* (*rhlABC*, *rhll*, and *rhlR*) and the *pqsABCDEH* systems (**Table S6**), which collectively orchestrate quorum sensing. In turn, quorum sensing coordinates diverse physiological processes, including biofilm formation, and regulates the production of various virulence factors [78]. To evaluate the impact of GCP disruption on virulence traits, assays were performed for measuring key virulence traits such as extracellular proteases, rhamnolipids, and pigments.

Production of the siderophore pyoverdine (**Fig. 7A**) and the phenazine pyocyanin (**Fig. 7B**) was significantly reduced in strain PA14Δ32, with levels < 13% and 1%, respectively, compared to the wild-type strain. Additionally, the mutant exhibited impaired rhamnolipid production (**Fig. 7C**) and decreased extracellular protease (exoprotease) activity (**Fig. 7D**), with reductions of ca. 80% and 40%, respectively. These findings align with low expression of the *rhlABC* genes involved in rhamnolipid biosynthesis. Significant transcriptional changes were also observed in *phzABCDE* and *pchABCDEFG*, genes responsible for pyocyanin and pyoverdine synthesis (**Table S6** and **Data S1**). Reduced exoprotease production correlated with transcriptional downregulation of genes involved in secretion systems, particularly type III and type IV systems (**Fig. 7D**, **Table S6**, and **Data S1**).

**Fig. 7.**
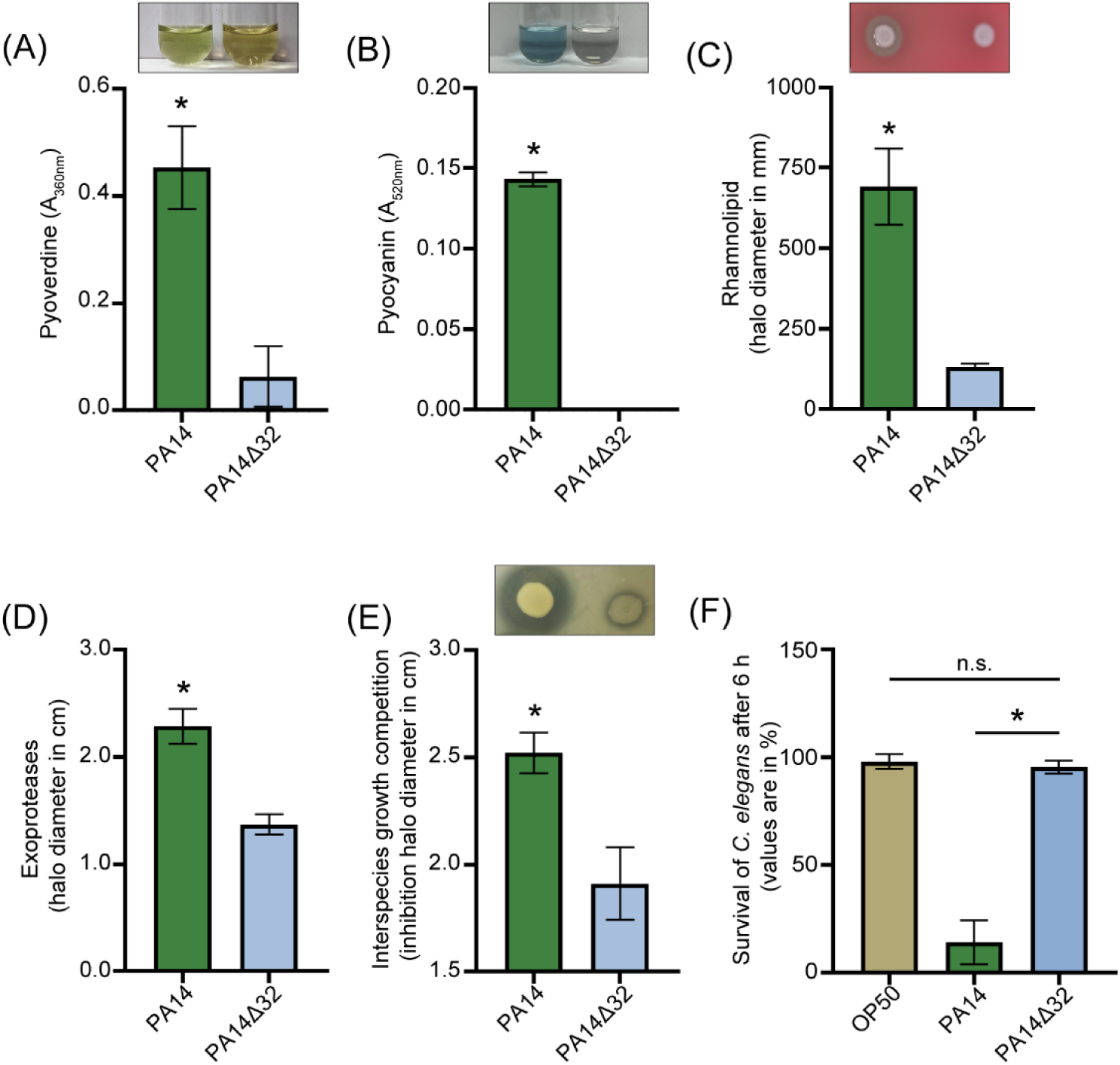
Virulence factor production in strain PA14Δ32. **(A)** Pyoverdine quantification from King’s B culture supernatants, measured by absorbance at 360 nm (A_360 nm_). **(B)** Pyocyanin extraction from LB culture supernatants using chloroform, quantified by absorbance at 520 nm (A_520 nm_). **(C)** Rhamnolipid production assessed *via* hemolysis on sheep blood agar; 5 μL of bacterial culture was spotted and incubated at 37°C for 96 h. **(D)** Protease activity determined by degradation zones on 1% (w/v) milk agar after incubation at 37°C overnight. **(E)** Competition between *P. aeruginosa* and *S. aureus* USA300 assessed on tryptic soy agar plates. *S. aureus* cultures (OD_600_ = 1) were spread, and a 5 μL spot of PA14 or PA14Δ32 culture was added; inhibition halos and colony diameters were measured after overnight incubation. **(F)** Paralysis assay with *C. elegans* exposed to *P. aeruginosa* strains grown on brain-heart infusion agar. Survival percentage was recorded after 6 h. Degradation halos and colony diameters in panels **(C)**, **(D)**, and **(E)** were analyzed using the ImageJ software. Statistical significance at *p* < 0.05 is marked by an asterisk (*) symbol; non-significant differences are denoted as *n*.*s*. (Student’s *t*-test). In all cases, results represent averages ± standard deviations triplicate or quadruplicate measurements from three independent experiments.

Antibiotic resistance is potentially influenced by c-di-GMP levels [79,80]. To investigate this aspect, we determined the minimum inhibitory concentration (MIC) of the PA14Δ32 mutant and compared it with the wild-type strain using a panel of clinically relevant antibiotics, including ceftazidime, amikacin, and gentamicin. *P*. *aeruginosa* PA14Δ32 exhibited resistance levels comparable to the wild-type strain across nearly all antibiotics tested (**Table S8**). These molecular insights into virulence were further examined in a dual bacterial infection model, as described below.

### 3.7. Reduced virulence and diminished competitiveness against *Staphylococcus aureus* following GCP disruption

In cystic fibrosis and chronically infected wounds, *P. aeruginosa* and *Staphylococcus aureus* often coexist as comorbid human pathogens. Pyocyanin-mediated inter- and intracellular signaling allows *P. aeruginosa* to outcompete *S. aureus*, providing a clear competitive advantage [81,82]. Additionally, *P. aeruginosa* may utilize exoproteases to compete with staphylococci and sequester iron from *S. aureus via* the iron-chelating molecule pyoverdine [81,83].

Given the observed impairment in extracellular virulence factors upon elimination of GGDEF-domain proteins (**Fig. 7** and **Table S6**), we investigated how altered c-di-GMP levels influence interspecies competition with *S. aureus*. In co-culture assays, *P*. *aeruginosa* PA14Δ32 exhibited a significant 1.3-fold reduction in inhibitory activity against *S*. *aureus* strain USA300 compared to the wild-type strain (**Fig. 7E**). These findings, once again, underscore the importance of c-di-GMP in modulating interspecies interactions, a key aspect of virulence. To further explore the pathogenicity profile of the PA14Δ32 mutant, the free-living nematode *Caenorhabditis elegans* was used as a model organism [84]. A paralysis assay was performed to compare the effects of the wild-type strain and the PA14Δ32 mutant on adult worms exposed to bacterial lawns grown on brain-heart agar. The PA14Δ32 mutant had markedly reduced virulence; ca. 98% of the worms survived after 6 h, similar to the negative control with *E*. *coli* OP50 cells (**Fig. 7F**). In contrast, only ca. 16% of worms survived exposure to the wild-type strain (**Fig. 7F**). Taken together, these results highlight the essential role of c-di-GMP in regulating virulence and pathogenesis in *P*. *aeruginosa*. The next set of experiments was designed to correlate these phenotypes with key GGDEF-domain proteins.

### 3.8. Genetic complementation of strain PA14Δ32 with specific GGDEF-domain proteins partially restores wild-type phenotypes

Two GGDEF-domain proteins, WspR and PA14_23130, were selected for complementation in both the wild-type strain and the PA14Δ32 mutant to investigate the broader effects of altered c-di-GMP levels beyond GCP-specific influences (**Fig. 8**). WspR, a key component of the *P*. *aeruginosa* wrinkly spreader phenotype (Wsp) chemosensory system (**Fig. 8A**), is one most active DGCs in this species [27]. This system detects surface-induced cell envelope stress, promoting a biofilm lifestyle *via* c-di-GMP signaling [85,86]. In the PA14 strain, PA14_23130 is homologous to PA3177 of *P*. *aeruginosa* PAO1, which encodes an active DGC associated with biofilm drug tolerance (**Table S4**). However, its role in attachment and biofilm formation remains unclear [87]. These proteins were chosen for their cytoplasmic localization as GGDEF-only enzymes and their involvement in diverse cellular processes (**Table S4**).

**Fig. 8.**
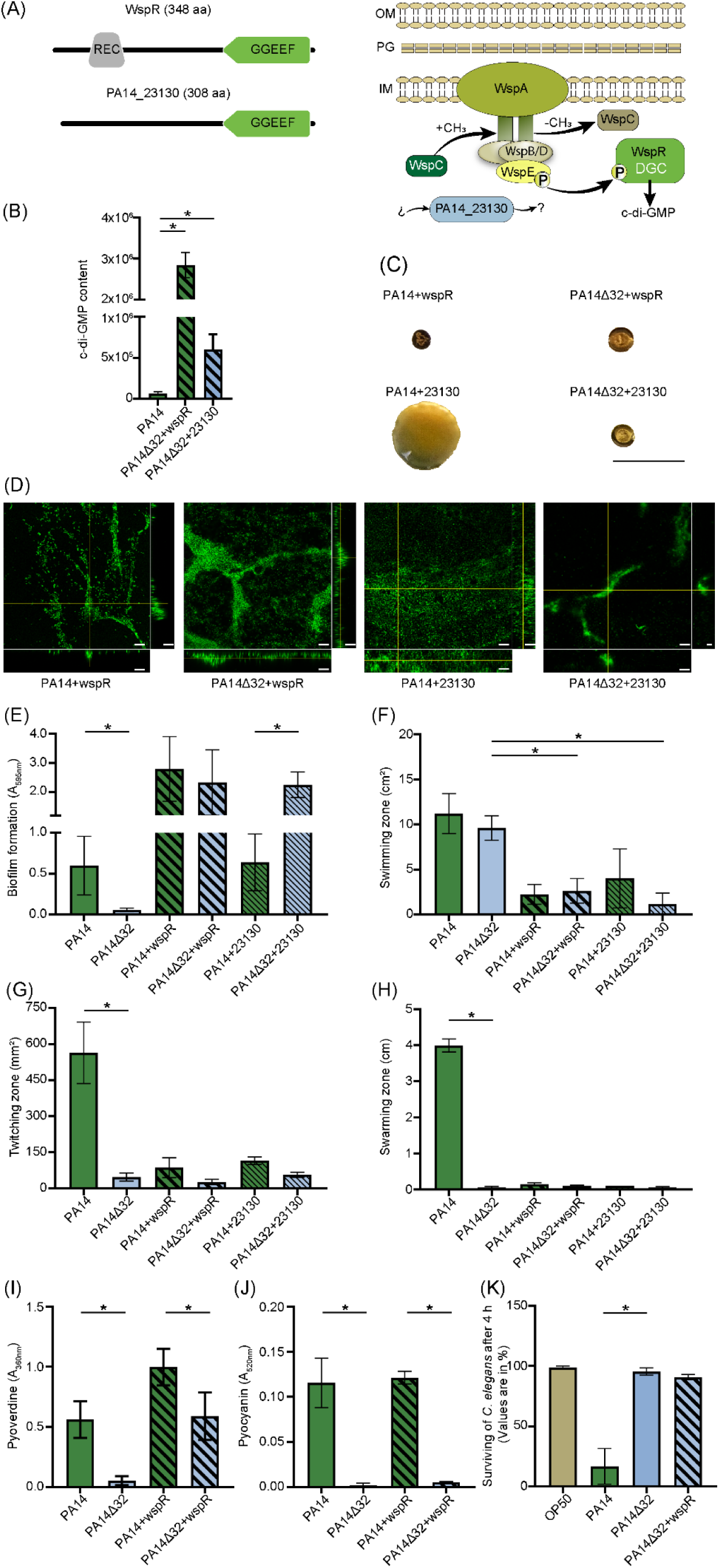
Complementation of strain PA14Δ32 with specific GGDEF-domain proteins partially restores wild-type phenotypes. **(A)** Domain architecture of the DGCs WspR and PA14_23130, both featuring the GGDEF domain, with WspR additionally containing a phospho-receiver (REC) domain. Both enzymes localize in the cytoplasm. The Wsp signal transduction complex comprises six proteins: WspA, an inner membrane (IM) receptor detecting surface-induced activation signals; WspE, which phosphorylates WspR, leading to c-di-GMP synthesis; WspC (methyltransferase) and WspF (methylesterase), which mediate signal adaptation; and WspB and WspD, scaffold proteins ensuring proper complex localization and function [119]. *Abbreviations*: *OM*, outer membrane; *PG*, peptidoglycan; and *GGDEF*, diguanylate cyclase domain. **(B)** LC-MS/MS analysis of c-di-GMP levels in PA14Δ32 strains expressing *wspR* (+ *wspR*) and *PA14_23130* (+ *23130*). Cultures were grown in LB supplemented with 0.1% (w/v) arabinose and normalized to the same OD_600_ before analysis. Results represent average c-di-GMP levels ± standard deviations. Characterization of **(C)** colony morphology (scale bar = 2 mm), **(D)** confocal laser scanning microscopy of biofilm structures in PA14 and PA14Δ32 strains transformed with plasmids pJN_*wspR* (+ *wspR*) or pJN_*23130* (+ *23130*), **(E)** biofilm formation, **(F)** swimming, **(G)** swarming, **(H)** twitching, **(I)** pyoverdine, **(J)** pyocyanin, and **(K)** *C*. *elegans* paralysis assays. Experimental conditions for each assay were consistent with those described in previous figures. For all complementation assays, strains PA14 and PA14Δ32 carried the empty plasmid pJN105 as an empty vector control. Measurements were performed in triplicate or quadruplicate across at least two independent experiments; results represent averages ± standard deviations. Statistical significance at *p* < 0.05 is indicated by an asterisk (*) symbol (one-way ANOVA with Tukey’s *post hoc* test).

Expression of *wspR* and *PA14_23130* in strain PA14Δ32 resulted in increased c-di-GMP content compared to the wild-type, with *wspR* expression producing the highest levels, supporting its role in c-di-GMP synthesis (**Fig. 8B**). Expression of *wspR* in PA14Δ32 and wild-type strains induced a small-colony variant (SCV) phenotype and fully restored biofilm formation in PA14Δ32 (**Fig. 8C** and **E**). Confocal microscopy confirmed the development of comparably structured, densely packed biofilms in both the wild type and PA14Δ32 expressing *wspR*, rescuing the biofilm deficiency in the multiple mutant (**Fig. 5C**). *PA14_23130* expression in the wild-type background did not affect biofilm formation, consistent with previous reports [87]. However, in the PA14Δ32 mutant, *PA14_23130* expression induced biofilm formation and SCV phenotypes (**Fig. 8C-E**).

To evaluate antibiotic resistance, the PA14Δ32 mutant engineered to express *PA14_23130* (referred to as PA14Δ32 + 23130) was tested against the same antibiotic panel described above (**Table S8**). Strain PA14Δ32 + 23130 showed resistance levels similar to the wild-type strain and the PA14Δ32 mutant for nearly all antibiotics tested, diverging from earlier reports. Notably, the absence of 31 GGDEF-domain proteins allowed this DGC to promote biofilm formation comparable to that observed with highly active *wspR* expression (**Fig. 8D** and **8E**).

While the analysis of motility produced the results anticipated (**Fig. 8F-H**), while pigment production followed an unexpected pattern. Expression of *wspR* in strain PA14Δ32 restored pyoverdine to wild-type levels (**Fig. 8I**), but pyocyanin production remained deficient (**Fig. 8J**), suggesting that other GGDEF-domain proteins with distinct functions are critical for this phenotype. The reduced virulence of the PA14Δ32 mutant, assessed *via C. elegans* paralysis assays, was unaffected by *wspR* expression (**Fig. 8K**), indicating that this DGC does not play a significant role in pathogenicity within this infection model.

## 4. DISCUSSION

In this study, we adopted a modular CRISPR/Cas9-based editing system [36] to disrupt the 32 GCP-encoding genes in *P*. *aeruginosa* PA14. This approach overcame previous challenges in genome editing for this bacterium, enabling the creation of a GCP-disrupted strain as a foundation for studying c-di-GMP signaling. By introducing premature *STOP* codons into these genes, we examined the broader functional consequences of GCP disruption, focusing on biofilm formation, motility, virulence, and antibiotic resistance. As a general conclusion, and despite disrupting all 32 ORFs, the PA14Δ32 mutant exhibited viability and growth patterns comparable to the wild-type strain across various carbon sources—indicating that c-di-GMP synthesis is nonessential for basic cellular functions.

The interplay between c-di-GMP and carbon metabolism in *P*. *aeruginosa* warrants further investigation, as studies in *E*. *coli*, *V*. *cholerae*, and *S*. *typhimurium* had revealed significant links between the second messenger and cell physiology. In *V*. *cholerae*, for instance, glucose-mediated interactions inactivate PdeS, increasing c-di-GMP levels and promoting biofilm formation [88]. Similarly, global regulators in *E*. *coli*, e.g., CsrA, influence both carbon metabolism and c-di-GMP-regulated traits [89]. In *S. enterica*, high levels of c-di-GMP repress the expression of genes involved in glucose (PTS^Glu^), mannose (PTS^Man^), and fructose (PTS^Fru^) uptake, guiding sugar metabolism to exopolysaccharide production [90]. Transcriptional analysis in our study highlighted downregulation of some genes, e.g., *oprB*, *gltK*, and ABC sugar transporter permeases, possibly linking c-di-GMP signaling to the uptake of glucose, glycerol, and fructose in *P*. *aeruginosa*. These findings pave the way for further research into the metabolic roles of c-di-GMP signaling in this pathogen.

The residual c-di-GMP levels detected in *P*. *aeruginosa* PA14Δ32 suggest the existence of additional synthesis pathways or cryptic functions encoding dinucleotide-processing functions [91–93], which underscores the complexity of c-di-GMP signaling characteristic of environmental bacteria. Complementation of the mutant with individual DGCs revealed phenotypes that highlight regulatory roles beyond c-di-GMP synthesis. For instance, reintroducing WspR in the mutant background partially restored biofilm formation and motility, suggesting that c-di-GMP levels alone do not dictate these traits and that specific GCP functions are needed for restoring wild-type–like phenotypes. These insights provide a framework to investigate individual DGC mechanisms in a system devoid of DGC redundancy, where the effect of single enzymes involved in c-di-GMP metabolism can be studied while avoiding crosstalk with similar functions. We previously applied a similar experimental framework by constructing a mutant of *P*. *putida* devoid of all major NADP^+^-dependent dehydrogenases [36], where native and heterologous enzymes producing the reducing nucleotide can be characterized in growth-coupled designs [94–96].

The model associating biofilm formation with reduced motility was supported by swimming behavior in experiments with the PA14Δ32 mutant, yet swarming and twitching deficiencies highlight additional complexity related to high c-di-GMP levels. Swarming, requiring flagellar function and rhamnolipid biosurfactants [50,97,98], is likely impacted by disrupted biosurfactant production in strain PA14Δ32. Previous studies implicated GGDEF-domain proteins in motility-related functions [99,100], suggesting their potential involvement in the regulation of swarming functions (including HsbD, NbdA, RoeA, SadC, BifA, GcbA, DipA, PA14_37690, and PA14_53310, all of which had altered transcription levels in the PA14Δ32 mutant). Comparative analyses with other bacteria, e.g., *Salmonella* [29] and *Caulobacter crescentus* [30], demonstrate diverse motility phenotypes in GCP-deficient strains (**Table S9**), emphasizing species-specific mechanisms. *D*. *zeae*, for instance, exhibited increased flagellum-mediated motility upon removal of GCPs [32], while motility was not affected in a similar mutant of *S*. *meliloti* [31].

To explore individual DGCs and their interactions, we employed an “all-but-one” disruption strategy, leaving only FimX intact in strain PA14Δ31. *P*. *aeruginosa* PA14Δ31 had normal twitching and exhibited properly assembled type IV pili. Indeed, introduction of FimX in the PA14Δ32 strain restored twitching and pili assembly, largely recapitulating the wild-type and PA14Δ31 phenotypes. Previous studies indicated that the degenerated GGDEF/EAL domains of FimX prevent c-di-GMP synthesis or degradation [27,101]. Instead, the PDE domain of this protein interacts with c-di-GMP, positively regulating pili assembly [102,103]. Our data further suggests that FimX can regulate twitching and pili assembly even in the absence of other GCPs—highlighting, once again, the value of a bacterial system devoid of DGC redundancy for this type of studies.

Transcriptomic analyses also revealed downregulation of quorum sensing and biofilm-related genes, reflecting an intricate relationship between c-di-GMP, quorum sensing, and virulence. Quorum sensing is intricately linked with signaling molecules, e.g., c-di-GMP and c-AMP, which mediate complex information translation in bacteria [78]. For instance, reduced c-di-GMP can upregulate the expression of the *rhl* and *pqs* genes, linked to regulators such as PqsR [104]. However, the correlation between c-di-GMP levels, GGDEF-domain proteins, and QS-mediated virulence factor expression remains poorly understood. In fact, comparing different cyclase-deficient strains (**Table S9**) reveals that the roles of c-di-GMP and DGCs in biofilm formation and virulence vary across bacteria. Each species has unique regulatory pathways and factors influencing c-di-GMP-related phenotypes [30,32,31,29]. This diversity challenges the notion of a universal mechanism and emphasizes the importance of considering species-specific traits in studying biofilm formation and virulence.

Our findings further exposed a reciprocal relationship between c-di-GMP levels and virulence, where elevated c-di-GMP levels suppress virulence factors—consistent with observations in *E*. *coli*, *S*. *enterica*, and *V*. *cholerae* [105,106,27,107,108]. The PA14Δ32 strain had reduced fitness in interspecies competition with *S*. *aureus*—two pathogens commonly coexisting and competing in clinical setups [109–111]. This phenotype could be linked to decreased production of anti-staphylococcal virulence factors, e.g., pyocyanin, a key virulence factor and a quorum sensing-signaling molecule [112]. FliA, the alternative σ factor required for flagellin synthesis, influences pyocyanin production *via* c-di-GMP and BifA [113]. Pyocyanin binds to RmcA, activating its phosphodiesterase activity [114], while BrlR functions as a receptor for both c-di-GMP and pyocyanin [115]. While *wspR* expression *in trans* restored pyoverdine levels in *P*. *aeruginosa* PA14Δ32, pyocyanin production remained significantly low. This observation indicates that merely increasing c-di-GMP levels alone is insufficient for pyocyanin synthesis, suggesting the involvement of additional DGCs beyond WspR. These findings also support the notion that c-di-GMP signaling operates in a localized or compartmentalized manner [116,117]. In the future, identifying the specific DGCs contributing to pyocyanin synthesis will contribute to our understanding of this intricate regulatory pathway—an effort supported by the availability of a bacterial system devoid of DGC redundancy.

The value of an “interference-free” bacterial system for dissecting individual enzymes or functions is illustrated by our experiments with *PA14_21130* expression. Poudyal and Sauer [87] highlighted a role for the PA3177 cyclase (similar to the PA14_23130 protein) as a c-di-GMP modulator that influences biofilm drug tolerance. Comparable biofilm architectures in the wild-type strain and the ΔPA3177 mutant indicated a negligible impact of PA3177 on biofilm formation [87]. Strempel et al. [118] proposed that PA3177 plays a role in the HClO-induced stress response, affecting motility, biofilm formation, and macrophages interaction. In our study, expressing *PA14_21130*, the orthologue of *PA3177* present in strain PA14, in the wild-type background did not affect biofilm production or architecture. However, *PA14_21130* expression in the “interference-free” PA14Δ32 background significantly influenced biofilm formation, suggesting a role for this DGC that could not be detected in the wild-type context. Hence, the genetic dissection approach adopted in our study offers a robust framework for addressing unresolved questions connected to the complex c-di-GMP signaling in *P*. *aeruginosa*, a model bacterial pathogen.

## Acknowledgments

The financial support provided by the Agencia Nacional de Promoción Científica y Tecnológica (ANPCyT) through grant PICT-2019-1590, the Consejo Nacional de Investigaciones Científicas y Técnicas (CONICET) through grant PIP-2022-11220210100945CO, and the Secretaría de Ciencia y Tecnología de la Universidad Nacional de Córdoba (SECYT-UNC) through grant 33620180100413CB to A.M.S. P.I.N. gratefully acknowledges financial support from The Novo Nordisk Foundation through grants NNF20CC0035580, *LiFe* (NNF18OC0034818), *TARGET* (NNF21OC0067996), FM·*Pseudomonas* (NNF24OC0091501), and *NovoF* (NNF23OC0083631), and the European Union’s Horizon2020 Research and Innovation Programme under grant agreements Nos. 814418 (*SinFonia*) and 101082049 (*TOLERATE*). R.A.M. received support from a postdoctoral fellowship from ANPCyT and Consejo Nacional de Ciencia y Tecnología (CONICET), as well as a Research and Training Grant from the Federation of European Microbiological Societies (FEMS). The responsibility for the content of this article rests solely with the authors, and the funding sources are not responsible for any use that may be made of the information contained therein.

## Data Availability

Data supporting the findings of this study can be found in the paper and its supplementary information. The WGS and the RNA-seq raw data generated in this study for the PA14Δ32 strain has been deposited in the public database GenBank under the BioProject ID PRJNA987335, which can be accessed at https://www.ncbi.nlm.nih.gov/bioproject/. Any additional data can be made available from the authors upon reasonable request.

## Supporting Information

Additional supporting information can be found online in the Supporting Information section.

