## Supplemental Data for "Genetic dissection of cyclic di-GMP signaling in *Pseudomonas aeruginosa via* diguanylate cyclase disruption"

by

Román A. Martino[1,2]†, Daniel C. Volke[3]†, Albano H. Tenaglia[1,2], Paula M. Tribelli[4,5], Pablo I. Nikel[3]* and Andrea M. Smania[1,2]*

[1] *Universidad Nacional de Córdoba, Facultad de Ciencias Químicas, Departamento de Química Biológica Ranwel Caputto, Córdoba, Argentina*
[2] *CONICET, Universidad Nacional de Córdoba, Centro de Investigaciones en Química Biológica de Córdoba (CIQUIBIC), Córdoba, Argentina*
[3] *The Novo Nordisk Foundation Center for Biosustainability, Technical University of Denmark, Kongens Lyngby, Denmark*
[4] *Universidad de Buenos Aires, Facultad de Ciencias Exactas y Naturales, Departamento de Química Biológica, Buenos Aires, Argentina*
[5] *CONICET, Universidad de Buenos Aires, Instituto de Química Biológica de la Facultad de Ciencias Exactas y Naturales (IQUIBICEN), Buenos Aires, Argentina*

† These authors contributed equally to this work.

---

* *Correspondence to:*

or

#### SUPPLEMENTARY FIGURES

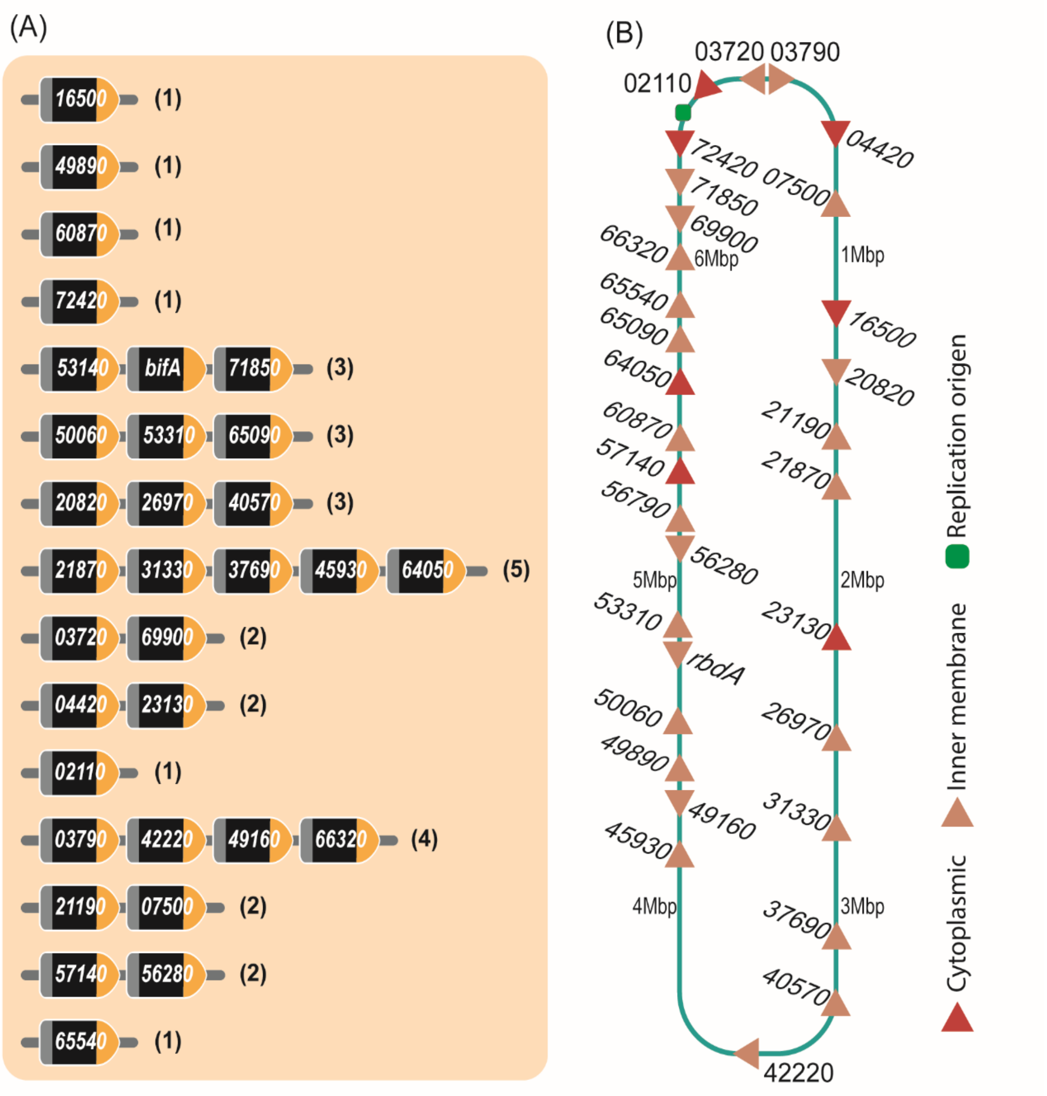


**Figure S1. (A)** Sequential interruption of the c-di-GMP signaling network in *Pseudomonas aeruginosa* PA14, resulting in the generation of strain PA14Δ32 and intermediate mutant strains. The number of targets per round is indicated between parentheses. **(B)** Physical map of the *P. aeruginosa* PA14 genome (not drawn to scale), showing the location of open reading frames (ORFs) encoding proteins with predicted GGDEF domains.

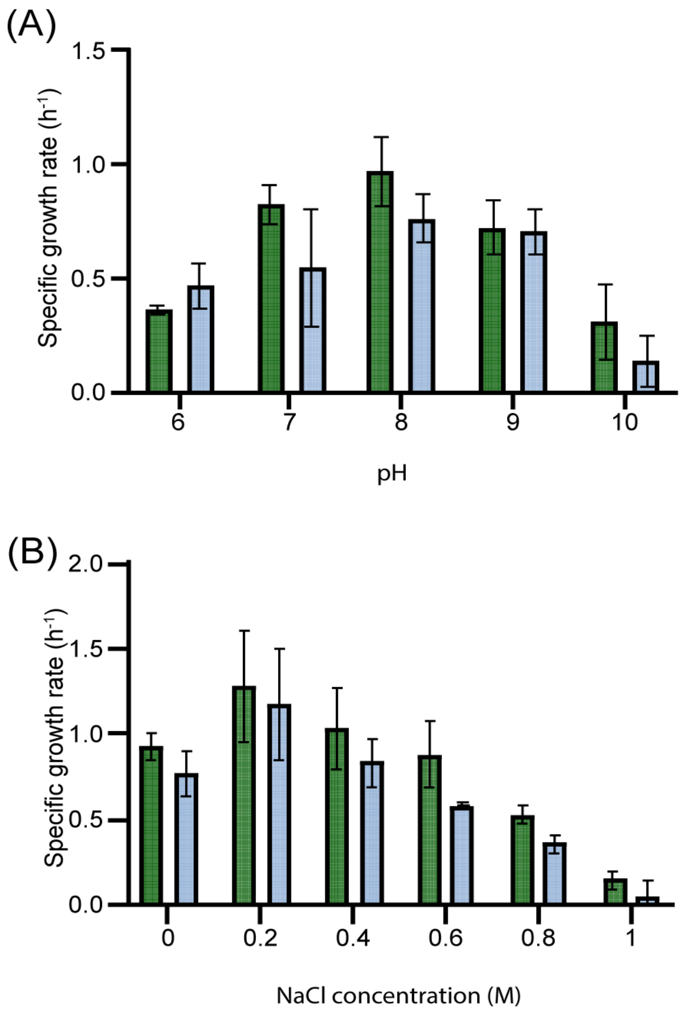


**Figure S2. (A)** and **(B)** Specific growth rate comparison between *P. aeruginosa* PA14 (parental strain, green bars) and strain PA14Δ32 (blue bars) under various stress conditions. Cultures were inoculated in LB adjusted to different pH levels and NaCl concentrations and incubated in a plate reader for 24 h at 37°C with agitation. The results represent averages standard deviation from biological triplicates.

### Supplementary Tables

**Table S1.** Bacterial strains and plasmids used in this study.

| **Strain** | **Relevant characteristics**[a] | **Reference** |
|---|---|---|
| *Escherichia coli* | | |
| DH5$\alpha$ $\lambda$*pir* | Cloning host; F$^{-}$ $\lambda^{-}$ *endA1 glnX44*(AS) *thiE1 recA1 relA1 spoT1 gyrA96*(Nal$^{R}$) *rfbC1 deoR nupG* $\Phi$80(*lacZ*$\Delta$*M15*) $\Delta$(*argF-lac*) *U169 hsdR17*($r_K^-$ $m_K^+$), $\lambda$*pir* lysogen | Platt et al. [1] |
| HB101 | Helper strain; F$^{-}$ $\lambda^{-}$ *hsdS20*($r_B^-$ $m_B^-$) *recA13 leuB6*(Am) *araC14* $\Delta$(*gpt-proA*)*62 lacY1 galK2*(Oc) *xyl-5 mtl-1 thiE1 rpsL20*(Sm$^{R}$) *glnX44*(AS) | Boyer and Roulland-Dussoix [2] |
| PIR2 | Cloning host; F$^{-}$ $\Delta$*lac169 rpoS*(Am) *robA1 creC510 hsdR514 endA recA1 uidA*($\Delta$MluI)::*pir* | Thermo Fisher Scientific |
| *Pseudomonas aeruginosa* | | |
| UCBPP-PA14 | Wild-type strain (referred to as PA14 throughout this study) | Liberati et al. [3] |
| PA14$\Delta$31 | Derivative of strain PA14 with an inactivation (premature *STOP* codon) in *wspR*, *yfiN*, *morA*, PA14_72420, *rbdA*, *bifA*, PA14_71850, PA14_50060, PA14_53310, PA14_65090, PA14_20820, PA14_26970, PA14_40570, PA14_21870, PA14_31330, PA14_37690, PA14_45930, PA14_65540, PA14_03720, PA14_69900, PA14_04420, PA14_23130, PA14_02110, PA14_03790, PA14_42220, PA14_49160, PA14_66320, PA14_21190, and PA14_07500 | This work |
| PA14$\Delta$32 | Derivative of strain PA14 with all diguanylate cyclases inactivated by premature *STOP* codons (*wspR*, *yfiN*, *morA*, PA14_72420, *rbdA*, *bifA*, PA14_71850, PA14_50060, PA14_53310, PA14_65090, PA14_20820, PA14_26970, PA14_40570, PA14_21870, PA14_31330, PA14_37690, PA14_45930, PA14_65540, PA14_03720, PA14_69900, PA14_04420, PA14_23130, PA14_02110, PA14_03790, PA14_42220, PA14_49160, PA14_66320, PA14_21190, PA14_07500, PA14_57140, PA14_56280, and PA14_64050) | This work |
| **Plasmid**[b] | **Relevant characteristics**[a] | **Reference** |
| pRK600 | Cm$^{R}$; *oriV*(ColE1), *tra*$^{+}$ *mob*$^{+}$ functions from plasmid RK2 | Keen et al. [4] |
| pTnS1 | Amp$^{R}$; *ori*(R6K), TnsABCD transposase functions | Choi et al. [5] |
| pBG·PelA | Km$^{R}$ Gm$^{R}$; plasmid pBG [6] carrying the $P_{pelA}$ promoter as a $P_{pelA}$(BDC2)→*msfGFP* transcriptional fusion | Benedetti et al. [7] |
| pBG·PelA·Sm | Km$^{R}$ Sm$^{R}$; derivative of plasmid pBG·PelA with a Sm resistance cassette | This work |
| pJN105 | Gm$^{R}$; arabinose-inducible expression vector; $P_{araBAD}$ promoter; *araC*; *oriV*(pBBR1) | Newman and Fuqua [8] |
| pJN_*wspR* | Gm$^{R}$; derivative of vector pJN105 bearing a copy of *wspR* (*PA14_16500*) | This work |
| pJN_*23130* | Gm$^{R}$; derivative of vector pJN105 bearing a copy of *PA14_23130* | This work |
| pMBLe | Gm$^{R}$; IPTG inducible expression vector; *lacI*; $P_{tac}$ promoter, *lac* operator; *oriV*(pBBR1) | Colque et al. [9] |
| pMBLe_*fimX* | Gm$^{R}$; derivative of vector pMBLe bearing a copy of *fimX* (*PA14_65540*) | This work |
| pBEC6 | Gm$^{R}$; cytosine base editing plasmid; *oriV*(pRO1600/ColE1); *sacB* | Volke et al. [10] |
| pBEC8 | Ap$^{R}$; cytosine base editing plasmid; *oriV*(pRO1600/ColE1); *sacB* | Volke et al. [10] |
| pMBEC6 | Gm$^{R}$; multiplex cytosine base editing plasmid; *oriV*(pRO1600/ColE1); *sacB* | Volke et al. [10] |
| pMBEC8 | Ap$^{R}$; multiplex cytosine base editing plasmid; *oriV*(pRO1600/ColE1); *sacB* | Volke et al. [10] |
| pMBEC4 | Sm$^{R}$; multiplex cytosine base editing plasmid; *oriV*(pRO1600/ColE1); *sacB* | Volke et al. [10] |
| pBEC6·*wspR* | Gm$^{R}$; derivative of vector pBEC6 bearing the *wspR* spacer | This work |

| | | |
|---|---|---|
| pBEC8·*yfiN* | Ap$^R$; derivative of vector pBEC8 bearing the *yfiN* spacer | This work |
| pBEC6·*morA* | Gm$^R$; derivative of vector pBEC6 bearing the *morA* spacer | This work |
| pBEC8·72420 | Ap$^R$; derivative of vector pBEC8 bearing the *PA14_72420* spacer | This work |
| pBEC6·71850 | Ap$^R$; derivative of vector pBEC8 bearing the *PA14_71850* spacer | This work |
| pBEC8·*bifA* | Ap$^R$; derivative of vector pBEC8 bearing the *bifA* spacer | This work |
| pBEC6·*rbdA* | Gm$^R$; derivative of vector pBEC6 bearing the *rbdA* spacer | This work |
| pMBEC6·Cluster_A | Gm$^R$; derivative of vector pMBEC6 bearing the multiplex guide RNA with *20820-26970-40570-07500-21190-64050* spacers | This work |
| pMBEC6·Cluster_B | Gm$^R$; derivative of vector pMBEC6 bearing the multiplex guide RNA with *50060-53310-56280-57140-65090* spacers | This work |
| pMBEC6·Cluster_C | Gm$^R$; derivative of vector pMBEC6 bearing the multiplex guide RNA with *02110-03790-04420-23130* spacers | This work |
| pMBEC8·Cluster_D | Ap$^R$; derivative of vector pMBEC8 bearing the multiplex guide RNA with 21870-31330-37690-45930-66320 spacers | This work |
| pMBEC4·Cluster_E | Sm$^R$; derivative of vector pMBEC4 bearing the multiplex guide RNA with *03720-42220-49160-69900* spacers | This work |
| pMBEC6·Cluster_A.1 | Gm$^R$; derivative of vector pMBEC6 bearing the multiplex guide RNA with *21190-64050-07500* spacers | This work |
| pMBEC6·Cluster_A.2 | Gm$^R$; derivative of vector pMBEC6 bearing the multiplex guide RNA with *21190-07500* spacers | This work |
| pMBEC6·Cluster_B.1 | Gm$^R$; derivative of vector pMBEC6 bearing the multiplex guide RNA with *57140-56280* spacers | This work |
| pMBEC6·Cluster_C.1 | Gm$^R$; derivative of vector pMBEC6 bearing the multiplex guide RNA with *02110-03790* spacers | This work |
| pMBEC6·Cluster_C.2 | Gm$^R$; derivative of vector pMBEC6 bearing the multiplex guide RNA with *04420-23130* spacers | This work |
| pMBEC6·Cluster_E.1 | Gm$^R$; derivative of vector pMBEC6 bearing the multiplex guide RNA with *42220-49160-66320* spacers | This work |
| pMBEC6·65540 | Gm$^R$; derivative of vector pMBEC6 bearing the multiplex guide RNA with *65540* spacer | This work |

[a] *Antibiotic markers*: *Amp*, ampicillin; *Ap*, apramycin; *Cm*, chloramphenicol; *Gm*, gentamicin; *Km*, kanamycin; *Nal*, nalidix acid; and *Sm*, streptomycin.

[b] Cluster_*X* (with *X* = A, B, C, D, or E) indicates a pMBEC plasmid bearing multiple spacers for different GCPs.

**Table S2.** Oligonucleotides and spacers used in this study.

| Oligonucleotide | Sequence (5'→3') | Use |
|---|---|---|
| *wspR*-sp-F | GTGGCTCCAGGACCTGGTGATGCC | Base-editing |
| *wspR*-sp-R | AAACGGCATCACCAGGTCCTGGAG | Base-editing |
| *yfiN*-sp-F | GTGGCGACGGCCAGAAGATCGGTT | Base-editing |
| *yfiN*-sp-R | AAACAACCGATCTTCTGGCCGTCG | Base-editing |
| *morA*-sp-F | GTGGGGAAGACCAATACGCCGGAC | Base-editing |
| *morA*-sp-R | AAACGTCCGGCGTATTGGTCTTCC | Base-editing |
| PA14_72420-sp-F | GTGGCAGTGGCCAGCGCGACTTCG | Base-editing |
| PA14_72420-sp-R | AAACCGGGAAATCTGGCTCGACAG | Base-editing |
| PA14_71850-sp-F | GTGGGGTCCAGCATTCCTATCTCG | Base-editing |
| PA14_71850-sp-R | AAACCGAGATAGGAATGCTGGACC | Base-editing |
| *bifA*-sp-F | GTGGCACCAGTTGCCGCTGCTCAA | Base-editing |
| *bifA*-sp-R | AAACTTGAGCAGCGGCAACTGGTG | Base-editing |
| *rbdA*-sp-F | GTGGCGTCCAGTTGGTCATCTGCT | Base-editing |
| *rbdA*-sp-R | AAACAGCAGATGACCAACTGGACG | Base-editing |
| PA14_07500-F | ATCGAGGTCTCCGTCCGAACAGAAGTTCGCTTGTTTTAGAGCTAGAAATAGC | Construction of plasmid pMBEC·Cluster_A |
| PA14_07500-R | ATCGAGGTCTCCTCGGTTTCTTAGCTGCCTATACGG | |
| PA14_20820-F | ATCGAGGTCTCCGTGGCCGCCCACCACTCGAAACCCGTTTTAGAGCTAGAAATAGC | |
| PA14_20820-R | ATCGAGGTCTCCGCGCTTTCTTAGCTGCCTATACGG | |
| PA14_21190-F | ATCGAGGTCTCCCCGACCCATTGCCAGCGGCGGTTTTAGAGCTAGAAATAGC | |
| PA14_21190_64050-R | ATCGAGGTCTCCAAACGTGAAGAATTCCTGGCCGCCTTTCTTAGCTGCCTATACGG | |
| PA14_26970-F | ATCGAGGTCTCCGCGCGGCCAGGTGGATGGCTGTTTTAGAGCTAGAAATAGC | |
| PA14_26970-R | ATCGAGGTCTCCGGGTTTTCTTAGCTGCCTATACGG | |
| PA14_40570-F | ATCGAGGTCTCCACCCAACTGGGCGCGACCCTGTTTTAGAGCTAGAAATAGC | |
| PA14_40570-R | ATCGAGGTCTCCGGACTTTCTTAGCTGCCTATACGG | |
| PA14_50060-F | ATCGAGGTCTCCGTGGTGCGCCACAGCTTGGGTGGCGTTTTAGAGCTAGAAATAGC | Construction of plasmid pMBEC·Cluster_B |
| PA14_50060-R | ATCGAGGTCTCCCGGGTTTCTTAGCTGCCTATACGG | |
| PA14_53310-F | ATCGAGGTCTCCCCCGACCAGCGCGTGCTCTTGTTTTAGAGCTAGAAATAGC | |
| PA14_53310-R | ATCGAGGTCTCCCATCTTTCTTAGCTGCCTATACGG | |
| PA14_56280-F | ATCGAGGTCTCCGATGCGCCAACTGGAAGACCGTTTTAGAGCTAGAAATAGC | |
| PA14_56280-R | ATCGAGGTCTCCGCAGTTTCTTAGCTGCCTATACGG | |
| PA14_57140-F | ATCGAGGTCTCCCTGCGCCAGGTGGAAACCCGGTTTTAGAGCTAGAAATAGC | |
| PA14_57140_65090-R | ATCGAGGTCTCCAAACCAGTCGCAGCGTCTGGTGGCTTTCTTAGCTGCCTATACGG | |
| PA14_02110-F | ATCGAGGTCTCCGTGGCCACGGCCAGAGCTACGTGCGTTTTAGAGCTAGAAATAGC | Construction of plasmid pMBEC·Cluster_C |
| PA14_02110-R | ATCGAGGTCTCCGCCGTTTCTTAGCTGCCTATACGG | |
| PA14_03790-F | ATCGAGGTCTCCCGGCCACCAGGTCGGCGACCGTTTTAGAGCTAGAAATAGC | |
| PA14_03790-R | ATCGAGGTCTCCGGCCTTTCTTAGCTGCCTATACGG | |
| PA14_04420-F | ATCGAGGTCTCCGGCCAGTGGCCTCTACCGAAGTTTTAGAGCTAGAAATAGC | |
| PA14_04420-R | ATCGAGGTCTCCGACTTTTCTTAGCTGCCTATACGG | |
| PA14_23130-F | ATCGAGGTCTCCAGTCCGCCAACTGATCCCGCGTTTTAGAGCTAGAAATAGC | |
| PA14_23130_53140-R | ATCGAGGTCTCCAAACAGCAGATGACCAACTGGACGTTTCTTAGCTGCCTATACGG | |

| | | |
|---|---|---|
| PA14_21870-F | ATCGAGGTCTCCGTGGGGCCCGAGCCTCTCGCGCCCGTTTTAGAGCTAGAAATAGC | Construction of plasmid pMBEC·Cluster_D |
| PA14_21870-R | ATCGAGGTCTCCTTGCTTTCTTAGCTGCCTATACGG | |
| PA14_31330-F | ATCGAGGTCTCCGCAACAGGAAGAAACCGAGTGTTTTAGAGCTAGAAATAGC | |
| PA14_31330-R | ATCGAGGTCTCCAGTCTTTCTTAGCTGCCTATACGG | |
| PA14_37690-F | ATCGAGGTCTCCGACTTCCCAGATCCAGTCCGGTTTTAGAGCTAGAAATAGC | |
| PA14_37690-R | ATCGAGGTCTCCAGGCTTTCTTAGCTGCCTATACGG | |
| PA14_45930-F | ATCGAGGTCTCCGCCTACCAGGACAGTCTCACGTTTTAGAGCTAGAAATAGC | |
| PA14_45930-R | ATCGAGGTCTCCTGCATTTCTTAGCTGCCTATACGG | |
| PA14_03720-F | ATCGAGGTCTCCGTGGGCCAACTGACCCACTACGTCGTTTTAGAGCTAGAAATAGC | |
| PA14_03720-R | ATCGAGGTCTCCCGGCTTTCTTAGCTGCCTATACGG | |
| PA14_12810-F | ATCGAGGTCTCCCTACCAGCCCAAGGTGGCCCGTTTTAGAGCTAGAAATAGC | Construction of plasmid pMBEC·Cluster_E |
| PA14_12810_14530-R | ATCGAGGTCTCCAAACGCTCGGACGTCTCCTGGAGGTTTCTTAGCTGCCTATACGG | |
| PA14_42220 -F | ATCGAGGTCTCCGCCGGCCGAGGCATCGACAAGTTTTAGAGCTAGAAATAGC | |
| PA14_42220 -R | ATCGAGGTCTCCGGTATTTCTTAGCTGCCTATACGG | |
| PA14_49160-F | ATCGAGGTCTCCTACCAGCGCGCCCACATTTCGTTTTAGAGCTAGAAATAGC | |
| PA14_49160-R | ATCGAGGTCTCCTGCGTTTCTTAGCTGCCTATACGG | |
| PA14_69900-F | ATCGAGGTCTCCCGCATGCCACATCAGCGGATGTTTTAGAGCTAGAAATAGC | |
| PA14_69900-R | ATCGAGGTCTCCGTAGTTTCTTAGCTGCCTATACGG | |
| PA14_07500-3-F | ATCGAGGTCTCCCGAACAGCACTACCGGGGACGTTTTAGAGCTAGAAATAGC | |
| PA14_07500-3_07500-2-R | ATCGAGGTCTCCAAACCCGACCCAGGGGGCCTGGAGTTTCTTAGCTGCCTATACGG | |
| PA14_21190-3-F | ATCGAGGTCTCCGTGGCTGCACCAGTCCAAGGCCTGGTTTTAGAGCTAGAAATAGC | |
| PA14_21190-3-R | ATCGAGGTCTCCCGCATTTCTTAGCTGCCTATACGG | |
| PA14_21190-4-F | ATCGAGGTCTCCTGCGGCCCAGATGCCGCAGGGTTTTAGAGCTAGAAATAGC | Construction of plasmid pMBEC·Cluster_A.1 |
| PA14_21190-4-R | ATCGAGGTCTCCCGCCTTTCTTAGCTGCCTATACGG | |
| PA14_64050-2-F | ATCGAGGTCTCCGGCGGCCAGGAATTCTTCACGTTTTAGAGCTAGAAATAGC | |
| PA14_64050-2-R | ATCGAGGTCTCCGCTTTTTCTTAGCTGCCTATACGG | |
| PA14_64050-3-F | ATCGAGGTCTCCAAGCAGGTCAACGACCGTTTGTTTTAGAGCTAGAAATAGC | |
| PA14_64050-3-R | ATCGAGGTCTCCTTCGTTTCTTAGCTGCCTATACGG | |
| PA14_56280-2-F | ATCGAGGTCTCCCTTGCCAACTGGCCTTCCTCGTTTTAGAGCTAGAAATAGC | |
| PA14_56280-2_56280-3-R | ATCGAGGTCTCCAAACGCGGCGGCAGTTGGAACACGTTTCTTAGCTGCCTATACGG | |
| PA14_57140-2-F | ATCGAGGTCTCCGTGGGCAACAGCTGGTCCGGCCGCGTTTTAGAGCTAGAAATAGC | Construction of plasmid pMBEC·Cluster_B.1 |
| PA14_57140-2-R | ATCGAGGTCTCCGGCGTTTCTTAGCTGCCTATACGG | |
| PA14_57140-3-F | ATCGAGGTCTCCCGCCAGGTGGAAACCCGCGGGTTTTAGAGCTAGAAATAGC | |
| PA14_57140-3-R | ATCGAGGTCTCCGAGGTTTCTTAGCTGCCTATACGG | |
| PA14_57140-4-F | ATCGAGGTCTCCCCTCAGCCAGGCCGGCTACCGTTTTAGAGCTAGAAATAGC | |
| PA14_57140-4-R | ATCGAGGTCTCCCAAGTTTCTTAGCTGCCTATACGG | |
| PA14_02110-2-F | ATCGAGGTCTCCGTGGGCCCAACCGACGCATGCTGCGTTTTAGAGCTAGAAATAGC | |
| PA14_02110-2-R | ATCGAGGTCTCCGGCGTTTCTTAGCTGCCTATACGG | |

| | | |
|---|---|---|
| PA14_02110-3-F | ATCGAGGTCTCCCGCCGCCCCAGCGGCCGCAGGTTTTAGAGCTAGAAATAGC | Construction of plasmid pMBEC·Cluster_C.1 |
| PA14_02110-3-R | ATCGAGGTCTCCGGAGTTTCTTAGCTGCCTATACGG | |
| PA14_03790-2-F | ATCGAGGTCTCCCTCCAGCGTATCGCCCAGAGGTTTTAGAGCTAGAAATAGC | |
| PA14_03790-2_03790-3-R | ATCGAGGTCTCCAAACGCCTCGCCCTGGCGACTGACTTTCTTAGCTGCCTATACGG | |
| PA14_42220-2-F | ATCGAGGTCTCCGTGGCGACCAGATGCCGAAGCCCAGTTTTAGAGCTAGAAATAGC | |
| PA14_42220-2-R | ATCGAGGTCTCCGTACTTTCTTAGCTGCCTATACGG | |
| PA14_42220-3-F | ATCGAGGTCTCCGTACCAGATATCCCGCCACGGTTTTAGAGCTAGAAATAGC | Construction of plasmid pMBEC·Cluster_E.1 |
| PA14_42220-3-R | ATCGAGGTCTCCTGGATTTCTTAGCTGCCTATACGG | |
| PA14_49160-2-F | ATCGAGGTCTCCTCCAGAACGTCACCAAGGCCGTTTTAGAGCTAGAAATAGC | |
| PA14_49160-2-R | ATCGAGGTCTCCGAACTTTCTTAGCTGCCTATACGG | |
| PA14_49160-3-F | ATCGAGGTCTCCGTTCCAGGAGATCACCTATGGTTTTAGAGCTAGAAATAGC | |
| PA14_49160-3-R | ATCGAGGTCTCCGGGTTTTCTTAGCTGCCTATACGG | |
| PA14_66320-2-F | ATCGAGGTCTCCACCCAGCAGAACAGCTGGCAGTTTTAGAGCTAGAAATAGC | |
| PA14_66320-2-R | ATCGAGGTCTCCGCAGTTTCTTAGCTGCCTATACGG | |
| PA14_66320-3-F | ATCGAGGTCTCCCTGCCACATGCGGTTCTCGCGTTTTAGAGCTAGAAATAGC | |
| PA14_66320-3-R-AAC | ATCGAGGTCTCCAAACTTTCTTAGCTGCCTATACGG | |
| PA14_07500-4-F | ATCGAGGTCTCCTTTCCCAGGCAATGGCATTCGTTTTAGAGCTAGAAATAGC | |
| PA14_07500-4-R | ATCGAGGTCTCCAAACTTTCTTAGCTGCCTATACGG | |
| PA14_07500-5-F | ATCGAGGTCTCCGGAACCAGTTGGAACGCGACGTTTTAGAGCTAGAAATAGC | Construction of plasmid pMBEC·Cluster_A.2 |
| PA14_07500-5-R | ATCGAGGTCTCCCAGGTTTCTTAGCTGCCTATACGG | |
| PA14_07500-11-F | ATCGAGGTCTCCCGACCAGACACCGGCGCTGGGTTTTAGAGCTAGAAATAGC | |
| PA14_07500-11-R | ATCGAGGTCTCCTGCATTTCTTAGCTGCCTATACGG | |
| PA14_21190-4-F | ATCGAGGTCTCCCCTGCGGCATCTGGGCCGCAGTTTTAGAGCTAGAAATAGC | |
| PA14_21190-4-R | ATCGAGGTCTCCGAAATTTCTTAGCTGCCTATACGG | |
| PA14_21190-7-F | ATCGAGGTCTCCTGCACCAGTCCAAGGCCTGCGTTTTAGAGCTAGAAATAGC | |
| PA14_21190-7-R | ATCGAGGTCTCCTTCCTTTCTTAGCTGCCTATACGG | |
| PA14_21190-9-F | ATCGAGGTCTCCGTGGCCCATTGCCAGCGGCGCGGGGTTTTAGAGCTAGAAATAGC | |
| PA14_21190-9-R | ATCGAGGTCTCCGTCGTTTCTTAGCTGCCTATACGG | |
| PA14_04420-2-F | ATCGAGGTCTCCGTGGTTCCAGGTCTCCACGCTCTCGTTTTAGAGCTAGAAATAGC | |
| PA14_04420-2-R | ATCGAGGTCTCCCGCCTTTCTTAGCTGCCTATACGG | |
| PA14_04420-3-F | ATCGAGGTCTCCGGCGCATCCATTCATGGTGCGTTTTAGAGCTAGAAATAGC | Construction of plasmid pMBEC·Cluster_C.2 |
| PA14_04420-3-R | ATCGAGGTCTCCTAGGTTTCTTAGCTGCCTATACGG | |
| PA14_23130-2-F | ATCGAGGTCTCCCCTACAGTTGCAGAGCAGCCGTTTTAGAGCTAGAAATAGC | |
| PA14_23130-2_23130-3-R | ATCGAGGTCTCCAAACGTTCCTCCACCGCCTGGCGCTTTCTTAGCTGCCTATACGG | |
| PA14_02110-seq-F | GCAACTGGTGCGGCTGGAGC | |
| PA14_02110-seq-R | GGAGGCTCAGCGCGCTGGAG | |
| PA14_65540-F | ATCGAGGTCTCCGTGGCAGGTCCCAGCTTTGCTGGAGTTTTAGAGCTAGAAATAGC | Construction of plasmid pMBEC6_65540 |
| PA14_65540-R | ATCGAGGTCTCCAAACTTTCTTAGCTGCCTATACGGCAGT | |
| PA14_03790-seq-F | CGGCATTCCCGAAGACCCGC | Sequencing |
| PA14_03790-seq-R | GCCCACGACGATGCAGTCCT | Sequencing |
| PA14_04420-seq-F | TCAGCGACGGGATCTGGGACTG | Sequencing |
| PA14_04420-seq-R | GGGTCTCGCCATCGTTCAGCAC | Sequencing |
| PA14_16500-seq-F | CCCCGGTCCCGGAGAGAAAC | Sequencing |
| PA14_16500-seq-R | ACGCGAGTGGTAGCGGATCC | Sequencing |

| | | |
|---|---|---|
| PA14_23130-seq-F | CTTGGCTTCGTAGAGCGCCTCG | Sequencing |
| PA14_23130-seq-R | ATTTCGACGCCGCCAAGCAGAA | Sequencing |
| PA14_26970-seq-F | GATTGCTCACCCGTAGCGCCTC | Sequencing |
| PA14_26970-seq-R | TGCTCGAACCGACCCACCAGAT | Sequencing |
| PA14_40570-seq-F | TGTTGCCCTCCTCGCTGTCGTA | Sequencing |
| PA14_40570-seq-R | GCCAGTTGCGCAACGGTTTTCG | Sequencing |
| PA14_49890-seq-F | ATACCGTCGAAGCCGCGGTG | Sequencing |
| PA14_49890-seq-R | GTCCTGCAAGCGCGCCTGCC | Sequencing |
| PA14_50060-seq-F | GTGGCCAGAACGACGCTCG | Sequencing |
| PA14_50060-seq-R | TTACCGCAGGCTTTCCGCGA | Sequencing |
| PA14_53310-seq-F | CGACGCCAGGCTCCCGGAA | Sequencing |
| PA14_53310-seq-R | CGCGCTACCCTGGTTGTTGC | Sequencing |
| PA14_56280-seq-F | GCGTTCGCAACAGCTTTCCAGC | Sequencing |
| PA14_56280-seq-R | AAAGCTCTTGCTCCGCCGGTTC | Sequencing |
| PA14_57140-seq-F | TACCGGCGCCCATCCTCTTCTC | Sequencing |
| PA14_57140-seq-R | GCATTCGTGCAGGTCCTCCACC | Sequencing |
| PA14_64050-seq-F | AGACCACGAAAGAAGCGCGTCA | Sequencing |
| PA14_64050-seq-R | AACTGAAGGTGCAGCGCAGGTC | Sequencing |
| PA14_65090-seq-F | CTGGGTATTGCCTGCCGTCG | Sequencing |
| PA14_65090-seq-R | CGAACAGCAGGTTGCCCTGG | Sequencing |
| PA14_72420-seq-F | GGAGGCAAGCAGGCCGTTC | Sequencing |
| PA14_72420-seq-R | GACGAGGCCTACGAACTGCCG | Sequencing |
| PA14_03720-seq-F | GCGTGTTGCATCAGGTGGTCGA | Sequencing |
| PA14_03720-seq-R | CTGCAGGGCCTGACCGACAATC | Sequencing |
| PA14_07500-seq-F | CCTGGGCGTACTTGAGGGTGGA | Sequencing |
| PA14_07500-seq-R | CTGATCCTGGTCTCGGTCGCCA | Sequencing |
| PA14_21190-seq-F | TACATCAGCGCGTCCTCGGTCA | Sequencing |
| PA14_21190-seq-R | TTGCAGGTGGCTCCCGCTATCT | Sequencing |
| PA14_21870-seq-F | GCATGAAGTCGTCACCGCCGAT | Sequencing |
| PA14_21870-seq-R | ACGTCTCTCCGGAGTCCCTCCT | Sequencing |
| PA14_31330-seq-F | GCCTTCTTCGGTTCCGATCGCC | Sequencing |
| PA14_31330-seq-R | TGGAAGAAGGTGCGGACGAGGT | Sequencing |
| PA14_37690-seq-F | CGTGCTGGTCGTTGATCGGCTT | Sequencing |
| PA14_37690-seq-R | GATCATGCGCCCATCGACCTGG | Sequencing |
| PA14_42220-seq-F | CGCCTTTTCCCTGATTGTGGCCA | Sequencing |
| PA14_42220-seq-R | AAGCAGTAGCCGTTCCGTCCCT | Sequencing |
| PA14_45930-seq-F | TACTTCGCCAGCATCCGGGTGA | Sequencing |
| PA14_45930-seq-R | GAGATAGGCGACCGGCGTGAGA | Sequencing |
| PA14_49160-seq-F | CGCGTTGCAGCTCCTCCTCGA | Sequencing |
| PA14_49160-seq-R | CTGTTCTTTCTCGCCGCCCTGG | Sequencing |
| PA14_53140-seq-F | CCAAGGCCTTCAGCGACGCC | Sequencing |
| PA14_53140-seq-R | GATACTCGAACTCGCGGCGG | Sequencing |
| PA14_56790-seq-F | CCTTCCACCCAGGCGGTCTACA | Sequencing |
| PA14_56790-seq-R | GGTCGCCGAGCTGGTAGGTGTA | Sequencing |
| PA14_60870-seq-F | AAAACGACCTCGAGCCAGGCTGGCTC | Sequencing |
| PA14_60870-seq-R | GCGCGAGCCAGCCTGGCTCGAGGTC | Sequencing |
| PA14_65540-seq-F | GAAGCCGAACGCCTGGTCAGTC | Sequencing |
| PA14_65540-seq-R | TACCAGAACGACCAGCGCCAGA | Sequencing |
| PA14_66320-seq-F | GCAGCGAACAGTTGCTCGACCT | Sequencing |
| PA14_66320-seq-R | GGGCCAGCTTGTTCTGCGTGAT | Sequencing |
| PA14_69900-seq-F | GCTGTCCTGGAACTGGCGGAAC | Sequencing |
| PA14_69900-seq-R | ACAGCTCTCCGAGGACAGCCTG | Sequencing |
| PA14_71850-seq-F | TACTCCACCCGGTACAGCTC | Sequencing |
| PA14_71850-seq-R | ATCCATCGTTTTCCTGCTGT | Sequencing |
| PA14_72420-seq-F | ATGCTGTTGACCCGCTCCAT | Sequencing |
| PA14_72420-seq-R | ACATCGATGCTCTCGGCG | Sequencing |

**Table S3.** Transitions and optimized parameters for c-di-GMP detection by LC-MS/MS.[a]

| **Compound** | **Parent mass** [m/z] | **Product mass** [m/z] | **DP** [V] | **EP** [V] | **CE** [V] | **CXP** [V] |
|---|---|---|---|---|---|---|
| c-di-GMP | 689 | 344 | –165 | –10 | –48 | –23 |
| c-di-GMP | 689 | 79 | –165 | –10 | –128 | –39 |
| c-di-GMP | 689 | 150 | –165 | –10 | –70 | –9 |

[a] *Abbreviations*: *DC*, declustering potential; *EP*, entrance potential; *CE*, collision energy; and *CXP*, collision cell exit potential.

**Table S4.** Open reading frames in *P. aeruginosa* PA14 encoding proteins with a GGDEF domain.[a]

| ORF | Name | Protein domain | Activity | Function | Gene length (bp) | Protein length (amino acids) | Reference(s) |
|---|---|---|---|---|---|---|---|
| PA14_02110 | *siaD* | GGEEF | DGC | Autoaggregation, biofilm formation, regulation of pyoverdine synthesis | 708 | 236 | [11-16] |
| PA14_03720 | — | GGDEF+ESL | PDE | Maintenance of basal levels of c-di-GMP | 2,283 | 761 | [17] |
| PA14_03790 | — | GGEEF | DGC | Maintenance of basal levels of c-di-GMP | 972 | 324 | [17] |
| PA14_04420 | — | GGEEF | N.D. | — | 1,131 | 377 | — |
| PA14_07500 | *rmcA* | GGDEF+EAL | PDE | Cellular redox sensing, L-arginine sensor, and biofilm maintenance | 3,738 | 1,246 | [18-20] |
| PA14_16500 | *wspR* | GGEEF | DGC | Biofilm formation | 1,044 | 348 | [21-27] |
| PA14_20820 | *hsbD* | GGEEF | DGC | Regulates twitching, swarming, chemotaxis, and biofilm formation | 1,170 | 390 | [28,22] |
| PA14_21190 | *nbdA* | AGDEF+EAL | PDE | Biofilm dispersal and motility (flagellar motor switching) | 2,358 | 786 | [29,30] |
| PA14_21870 | — | EAL+GGDDF | PDE | Biofilm formation *via* interaction with Pho regulon | 1,806 | 602 | [31,32] |
| PA14_23130 | — | GGEEF | DGC | Biofilm antimicrobial tolerance | 924 | 308 | [33,34] |
| PA14_26970 | — | GGEEF | DGC | — | 1,578 | 526 | [14] |
| PA14_31330 | — | SPTRF+EAL | PDE | Related to virulence factor expression and QS system in response to macrolides | 1,500 | 500 | [35,36] |
| PA14_37690 | — | GGDEF+EAL | PDE | Biofilm dispersal and swarming motility | 2,595 | 865 | [37-39] |
| PA14_40570 | — | GGEEF | DGC | — | 1,206 | 402 | [14] |
| PA14_42220 | *mucR* | GGDEF+EAL | DGC-PDE | Biofilm production, biofilm dispersal, surface attachment, and alginate production | 2,058 | 686 | [29,32,40-43] |
| PA14_45930[b] | *lapD* | RGGEF+KVL | DD | Surface attachment and biofilm formation | 1,953 | 651 | [44-51] |
| PA14_49160 | | GGDEF+EAL | DGC | Biofilm formation | 3,363 | 1,121 | [52-54] |
| PA14_49890 | *yfiN/ tpbB* | GGDEF | DGC | Overproduction of biofilm, resulting in a conversion to small and rugose colony variant phenotype, biofilm maintenance in response to oxidative stress | 1,308 | 436 | [22,55-63] |
| PA14_50060 | *roeA* | GGEEF | DGC | Biofilm formation, EPS production, and swarming motility | 1,197 | 399 | [22,63,64] |
| PA14_53140 | *rbdA* | GGDEF+EAL | PDE | Biofilm dispersal, EPS production, and motility (flagellar motor switching) | 2,472 | 824 | [65-68,30,37] |
| PA14_53310 | — | GGDEF | DGC | Motility | 2,046 | 682 | [14,22,69] |
| PA14_56280 | *sadC* | GGEEF | DGC | Gac/Rsm-mediated biofilm formation, EPS production, regulation of pyoverdine synthesis, and swarming motility | 1,128 | 376 | [14,12,15,16,63,70-76] |

| | | | | | | | |
|---|---|---|---|---|---|---|---|
| PA14_56790 | *bifA* | GGDQF+EAL | PDE | Biofilm formation, swarming motility, and maintenance of basal levels of c-di-GMP | 2,064 | 688 | [17,75,74,77,78] |
| PA14_57140 | | DEQHF | N.D. | — | 1,101 | 367 | — |
| PA14_60870 | *morA* | GGDEF+EAL | PDE | Biofilm formation and maintenance, virulence and type II secretion system regulation | 4,248 | 1,416 | [79-81] |
| PA14_64050 | *gcbA* | GGDDF | DGC | Initial surface adhesion for biofilm growth, biofilm dispersal, and swarming motility | 1,629 | 543 | [14,82-86] |
| PA14_65090 | *nicD* | GGDEF | DGC | Biofilm dispersal | 2,022 | 674 | [14,68,87] |
| PA14_65540 | *fimX* | GDSIF+EVL | PDE | Twitching motility | 2,076 | 692 | [37,88-96] |
| PA14_66320 | *dipA* | ASNEF+EAL | PDE | Biofilm dispersal, flagellum mediated swimming motility (flagellar motor switching), and maintenance of basal levels of c-di-GMP | 2,700 | 900 | [17,37,30,68,97-99] |
| PA14_69900 | *proE* | GSDEF+EAL | PDE | EPS production, conversion to small and rugose colony variant phenotype | 1,677 | 559 | [66] |
| PA14_71850 | — | AGDEF+EAL | N.D. | — | 2,853 | 951 | — |
| PA14_72420 | *dgcH* | GGEEF | DGC | Biofilm formation and maintenance of basal levels of c-di-GMP | 2,016 | 672 | [17] |

[a] Derived from homologous proteins in other *Pseudomonas* species, such as *P. fluorescens* Pf0-1 or *P. putida* KT2440.

[b] *Abbreviations*: *DD*, degenerate GGDEF and EAL/HD-GYP domains considered non-functional for c-di-GMP synthesis or degradation; *N.D.*, not experimentally determined.

**Table S5.** Spacers used to introduce premature *STOP* codons in the ORFs of strain PA14.

| ORF | Name | Protein domain | Gene length (bp) | Protein length (amino acids) | Spacer sequence (5'→3')[a] | Cytidine position[b] | Base before C | Mutation[c] |
|---|---|---|---|---|---|---|---|---|
| PA14_02110-2 | *siaD* | GGEEF | 708 | 236 | GCCCAACCGACGCATGCTGC | 8 | C | R71 |
| PA14_03720 | | GGDEF/ESL | 2,283 | 761 | GCCAACTGACCCACTACGTC | 3 | C | Q306 |
| PA14_03790-2 | | GGEEF | 972 | 324 | CTCCAGCGTATCGCCCAGAG | 4 | C | Q221 |
| PA14_04420-2 | | GGEEF | 1,131 | 377 | TTCCAGGTCTCCACGCTCTC | 4 | C | W99 |
| PA14_04420-3 | | GGEEF | 1,131 | 377 | GGCGCATCCATTCATGGTGC | 8 | T | W242 |
| PA14_07500-4 | *rmcA* | GGDEF/EAL | 3,738 | 1,246 | TTTCCCAGGCAATGGCATTC | 5-6 | C/C | W326 |
| PA14_07500-5 | *rmcA* | GGDEF/EAL | 3,738 | 1,246 | GGAACCAGTTGGAACGCGAC | 6 | C | Q747 |
| PA14_07500-11 | *rmcA* | GGDEF/EAL | 3,738 | 1,246 | CGACCAGACACCGGCGCTGG | 5 | C | Q536 |
| PA14_16500 | *wspR* | GGEEF | 1,044 | 348 | CTCCAGGACCTGGTGATGCC | 4 | C | Q69 |
| PA14_20820 | | GGEEF | 1,170 | 390 | CCGCCCACCACTCGAAACCC | 5-6 | C/C | W176 |
| PA14_21190-7 | | AGDEF/EAL | 2,358 | 786 | TGCACCAGTCCAAGGCCTGC | 6 | C | Q495 |
| PA14_21190-9 | | AGDEF/EAL | 2,358 | 786 | CCCATTGCCAGCGGCGCGGG | 2-3 | C/C | W121 |
| PA14_21870 | | EAL/GGDDF | 1,806 | 602 | GGCCCGAGCCTCTCGCGCCC | 5 | C | R222 |
| PA14_23130-2 | | GGEEF | 924 | 308 | CCTACAGTTGCAGAGCAGCC | 5-11 | A/G | Q45/Q47 |
| PA14_26970 | | GGEEF | 1,578 | 526 | GCGCGGCCAGGTGGATGGCT | 8 | C | Q309 |
| PA14_31330 | | SPTFR/EAL | 1,500 | 500 | GCAACAGGAAGAAACCGAGT | 2-5 | G/A | Q167/168 |
| PA14_37690 | | GGDEF/EAL | 2,595 | 865 | GACTTCCCAGATCCAGTCCG | 7-8 | C/C | W317 |
| PA14_40570 | | GGEEF | 1,206 | 402 | ACCCAACTGGGCGCGACCCT | 4 | C | Q143 |
| PA14_42220-2 | *mucR* | GDEF/EAL | 2,058 | 686 | CGACCAGATGCCGAAGCCCA | 4-5 | C/A | W56 |
| PA14_42220-3 | *mucR* | GDEF/EAL | 2,058 | 686 | GTACCAGATATCCCGCCACG | 5 | C | Q374 |
| PA14_45930 | *lapD* | RGGEF/KVL | 1,953 | 651 | GCCTACCAGGACAGTCTCAC | 7 | C | Q238 |
| PA14_49160-2 | | GGDEF/EAL | 3,363 | 1,121 | TCCAGAACGTCACCAAGGCC | 3 | C | Q669 |
| PA14_49160-3 | | GGDEF/EAL | 3,363 | 1,121 | GTTCCAGGAGATCACCTATG | 5 | C | Q346 |
| PA14_49890 | *yfiN/tpbB* | GGDEF | 1,308 | 436 | CGACGGCCAGAAGATCGGTT | 8 | C | Q138 |
| PA14_50060 | *roeA* | GGEEF | 1,197 | 399 | TGCGCCACAGCTTGGGTGGC | 5-6 | G/C | W96 |
| PA14_53140 | *rbdA* | GGDEF/EAL | 2,472 | 824 | CGTCCAGTTGGTCATCTGCT | 5 | C | W285 |
| PA14_53310 | | GGDEF | 2,046 | 682 | CCCGACCAGCGCGTGCTCTT | 7 | C | Q138 |
| PA14_56280-3 | *sadD* | GGDEF | 1,128 | 376 | CGTGTTCCAACTGCCGCCGC | 8 | C | Q122 |
| PA14_56790 | *bifA* | GGDQF/EAL | 2,064 | 688 | CACCAGTTGCCGCTGCTCAA | 4 | C | Q206 |
| PA14_57140-2 | | DEQHF | 1,101 | 367 | GCAACAGCTGGTCCGGCCGC | 5 | A | Q265 |

| | | | | | | | | |
|---|---|---|---|---|---|---|---|---|
| PA14_57140-3 | | DEQHF | 1,101 | 367 | CGCCAGGTGGAAACCCGCGG | 4 | C | Q222 |
| PA14_60870 | *morA* | GGDEF/EAL | 4,248 | 1,416 | GGAAGACCAATACGCCGGAC | 8 | C | Q201 |
| PA14_64050-2 | *gcbA* | GGDDF | 1,629 | 543 | GCCCAGCAACTGGAGTTCTT | 4-7 | C/G | Q141 |
| PA14_64050-3 | *gcbA* | GGDDF | 1,629 | 543 | AAGCAGGTCAACGACCGTTT | 4 | G | Q421 |
| PA14_64050-7 | *gcbA* | GGDDF | 1,629 | 1,629 | TGTTCCACTCCGCGCGGGTC | 5 | T | W40 |
| PA14_65090 | *nicD* | GGDEF | 2,022 | 674 | GCCACCAGACGCTGCGACTG | 5-6 | A/C | W120 |
| PA14_65540 | *fimX* | GDEF/EAL | 2,706 | 902 | CAGGTCCCAGCTTTGCTGGA | 7-8 | C-C | W53 |
| PA14_66320-3 | *dipA* | GDEF/EAL | 1,677 | 559 | CTGCCACATGCGGTTCTCGC | 4-5 | G/C | W314 |
| PA14_69900 | *proE* | GDEF/EAL | 1,677 | 559 | CGCATGCCACATCAGCGGAT | 8 | C | W262 |
| PA14_71850 | | GDEF/EAL | 2,853 | 951 | GGTCCAGCATTCCTATCTCG | 5 | C | Q301 |
| PA14_72420 | *dgcH* | GGDEF | 2,016 | 672 | CGACCAATCGCTGCGTGAGC | 5 | C | Q428 |

[a] The cytidine used as the substrate for the cytidine deaminase APOBEC, leading to the introduction of a premature *STOP* codon into the gene sequence, is indicated in red.

[b] Cytidine position from the 5'-spacer terminus.

[c] Amino acid residue and its position in the protein chain replaced by the premature *STOP* codon.

**Table S6.** Differentially expressed genes in strain PA14Δ32.

| Downregulated genes in strain PA14Δ32 compared to PA14 | | | |
|---|---|---|---|
| **ORF** | **$log_2$(fold-change)** | **Name** | **Product or function** |
| PA14_00560 | –2.4 | *exoT* | Exoenzyme T |
| PA14_00820 | –2.3 | *tagQ1* | Hypothetical protein |
| PA14_00850 | –2.0 | *tagS1* | ABC transporter permease |
| PA14_00860 | –2.4 | *tagT1* | ABC transporter ATP-binding protein |
| PA14_00875 | –2.3 | *ppkA* | Serine/threonine protein kinase PpkA |
| PA14_00890 | –2.2 | *pppA* | Serine/threonine-protein phosphatase |
| PA14_00910 | –2.2 | *icmF1* | Type VI secretion system membrane subunit TssM |
| PA14_00925 | –2.3 | *tssL1* | DotU family type VI secretion system protein |
| PA14_00940 | –2.0 | *tssK1* | Type VI secretion system baseplate subunit TssK |
| PA14_00960 | –2.1 | *tssJ1* | Type VI secretion system lipoprotein TssJ |
| PA14_01010 | –2.4 | *hsiB1* | Type VI secretion system contractile sheath small subunit |
| PA14_01020 | –2.4 | *hsiC1* | Type VI secretion system contractile sheath large subunit |
| PA14_01030 | –3.2 | *hcp1* | Type VI secretion system tube protein Hcp |
| PA14_01040 | –2.5 | *tagJ1* | Tetratricopeptide repeat protein |
| PA14_01060 | –3.0 | *tssE1* | Type VI secretion system baseplate subunit TssE |
| PA14_01070 | –3.1 | *tssF1* | Type VI secretion system baseplate subunit TssF |
| PA14_01080 | –3.0 | *tssG1* | Type VI secretion system baseplate subunit TssG |
| PA14_01100 | –2.2 | *clpV1* | Type VI secretion system ATPase TssH |
| PA14_01110 | –2.3 | *vgrG1a* | Type VI secretion system tip protein VgrG |
| PA14_01490 | –4.2 | | Hemolysin |
| PA14_01760 | –2.0 | *nuh* | Nonspecific ribonucleoside hydrolase |
| PA14_01780 | –2.4 | | Nucleoside 2-deoxyribosyltransferase |
| PA14_02230 | –2.4 | *cheW* | Purine-binding chemotaxis protein |
| PA14_02250 | –2.5 | *cheA* | Two-component sensor |
| PA14_06000 | –2.3 | | ClpA/B protease ATP binding subunit |
| PA14_09210 | –4.5 | *pchA* | Salicylate biosynthesis isochorismate synthase |
| PA14_09220 | –6.2 | *pchB* | Isochorismate-pyruvate lyase |
| PA14_09230 | –5.9 | *pchC* | Pyochelin biosynthetic protein PchC |
| PA14_09240 | –6.1 | *pchD* | Pyochelin biosynthesis protein PchD |
| PA14_09270 | –5.3 | *pchE* | Dihydroaeruginoic acid synthetase |
| PA14_09280 | –6.1 | *pchF* | Pyochelin synthetase |
| PA14_09290 | –6.1 | *pchG* | Pyochelin biosynthetic protein PchG |
| PA14_09300 | –5.4 | | ABC transporter ATP-binding protein |
| PA14_09320 | –7.2 | | ABC transporter ATP-binding protein |
| PA14_09340 | –5.6 | *fptA* | Fe(III)-pyochelin outer membrane receptor |
| PA14_09350 | –4.9 | | Hypothetical protein |
| PA14_09370 | –5.3 | | Hypothetical protein |
| PA14_09380 | –6.7 | | Transporter |
| PA14_09400 | –2.2 | *phzS* | Hypothetical protein |
| PA14_09440 | –2.6 | *phzE1* | Phenazine biosynthesis protein PhzE |
| PA14_09450 | –3.3 | *phzD1* | Phenazine biosynthesis protein PhzD |
| PA14_09460 | –3.9 | *phzC1* | Phenazine biosynthesis protein PhzC |

| | | | |
|---|---|---|---|
| PA14_09470 | –5.0 | *phzB1* | Phenazine biosynthesis protein |
| PA14_09480 | –8.4 | *phzA1* | Phenazine biosynthesis protein |
| PA14_09500 | –3.2 | *opmD* | Outer membrane protein |
| PA14_09520 | –2.7 | *mexI* | RND efflux transporter |
| PA14_09530 | –2.8 | *mexH* | RND efflux membrane fusion protein |
| PA14_09540 | –2.2 | *mexG* | Hypothetical protein |
| PA14_10350 | –2.2 | | Secretion protein |
| PA14_10360 | –3.6 | | Hypothetical protein |
| PA14_10380 | –3.1 | | Hypothetical protein |
| PA14_10490 | –3.4 | | Hypothetical protein |
| PA14_10500 | –4.2 | | *cbb3*-type cytochrome *c* oxidase subunit I |
| PA14_10530 | –2.2 | | GntR family transcriptional regulator |
| PA14_10540 | –5.5 | | Iron-sulfur cluster-binding protein |
| PA14_10550 | –5.6 | | Sulfite or nitrite reductas |
| PA14_10560 | –5.5 | | Hypothetical protein |
| PA14_11120 | –3.1 | | Response regulator |
| PA14_11810 | –3.3 | | Aldehyde dehydrogenase |
| PA14_13350 | –4.7 | | Hypothetical protein |
| PA14_13360 | –5.5 | | Hypothetical protein |
| PA14_13370 | –5.7 | | Hypothetical protein |
| PA14_13380 | –4.8 | | Hypothetical protein |
| PA14_13390 | –4.4 | | Hypothetical protein |
| PA14_16800 | –2.5 | | Efflux transmembrane protein |
| PA14_18630 | –4.4 | | Serine protease |
| PA14_18800 | –3.1 | | Hypothetical protein |
| PA14_18810 | –2.4 | | Hypothetical protein |
| PA14_19100 | –4.8 | *rhlA* | Rhamnosyltransferase chain A |
| PA14_19110 | –3.1 | *rhlB* | Rhamnosyltransferase chain B |
| PA14_19120 | –2.4 | *rhlR* | Transcriptional regulator RhlR |
| PA14_19130 | –5.1 | *rhlI* | Autoinducer synthesis protein RhlI |
| PA14_20610 | –3.9 | *lecB* | Fucose-binding lectin PA-IIL |
| PA14_20920 | –2.7 | | Hypothetical protein |
| PA14_20940 | –4.5 | | Acyl carrier protein |
| PA14_20950 | –5.9 | *fabH2* | 3-Oxoacyl-ACP synthase |
| PA14_20960 | –5.1 | | Isomerase |
| PA14_20970 | –5.4 | *cyp23* | Cytochrome P450 |
| PA14_20980 | –5.3 | | Short chain dehydrogenas |
| PA14_21000 | –6.0 | | Hypothetical protein |
| PA14_21010 | –6.4 | | FAD-dependent monooxygenase |
| PA14_21020 | –4.4 | | Non-ribosomal peptide synthetase |
| PA14_21030 | –3.9 | | ATP-dependent Clp protease proteolytic subunit |
| PA14_21190 | –2.0 | | Hypothetical protein |
| PA14_22980 | –3.6 | | Sugar ABC transporter substrate-binding protein |
| PA14_22990 | –3.9 | | ABC sugar transporter permease |
| PA14_23000 | –3.7 | | ABC sugar transporter permease |
| PA14_23010 | –4.1 | *gltK* | ABC transporter ATP-binding protein |

| | | | |
|---|---|---|---|
| PA14_23030 | –3.6 | *oprB* | Glucose/carbohydrate outer membrane porin OprB precursor |
| PA14_28050 | –2.5 | | Chemotaxis transducer |
| PA14_28360 | –3.0 | | Hypothetical protein |
| PA14_30560 | –4.9 | | Hypothetical protein |
| PA14_30570 | –4.0 | | Periplasmic spermidine/putrescine-binding protein |
| PA14_30580 | –4.1 | | LuxR family transcriptional regulator |
| PA14_30620 | –2.2 | | AraC family transcriptional regulator |
| PA14_30630 | –3.2 | *pqsH* | FAD-dependent monooxygenase |
| PA14_31450 | –2.4 | | Hypothetical protein |
| PA14_31970 | –2.1 | *czcC* | CzcC family cobalt/zinc/cadmium efflux transporter outer membrane protein |
| PA14_32140 | –4.2 | *antC* | Anthranilate dioxygenase reductase |
| PA14_32150 | –4.2 | *antB* | Anthranilate dioxygenase small subunit |
| PA14_32160 | –4.2 | *antA* | Anthranilate dioxygenase large subunit |
| PA14_32190 | –2.2 | *antR* | Transcriptional regulator |
| PA14_33290 | –2.0 | | Hypothetical protein |
| PA14_33560 | –2.0 | | Adhesion protein |
| PA14_33580 | –2.0 | | Hypothetical protein |
| PA14_33830 | –2.4 | | Hypothetical protein |
| PA14_34800 | –2.9 | | Amino acid transporter LysE |
| PA14_34810 | –3.9 | | Non-ribosomal peptide synthetase |
| PA14_34820 | –4.3 | *ambD* | Regulatory protein |
| PA14_34830 | –4.9 | *ambC* | Regulatory protein |
| PA14_34840 | –3.3 | | Non-ribosomal peptide synthetase |
| PA14_35160 | –2.1 | | Phenazine-utilizing monooxygenase A |
| PA14_36310 | –5.0 | *hcnC* | Hydrogen cyanide synthase HcnC |
| PA14_36320 | –5.7 | *hcnB* | Hydrogen cyanide synthase HcnB |
| PA14_36330 | –5.5 | *hcnA* | Hydrogen cyanide synthase HcnA |
| PA14_37745 | –2.2 | | Carbamoyl transferase |
| PA14_39970 | –3.7 | *phzA2* | Phenazine biosynthesis protein |
| PA14_39990 | –2.7 | | Hypothetical protein |
| PA14_40030 | –2.1 | | Hypothetical protein |
| PA14_40300 | –2.7 | | Hypothetical protein |
| PA14_40310 | –3.8 | | Acyl carrier protein |
| PA14_42300 | –2.5 | *pscG* | Type III export protein PscG |
| PA14_42310 | –2.2 | *pscF* | Type III export protein PscF |
| PA14_42360 | –2.3 | *pscB* | Type III export apparatus protein |
| PA14_42400 | –2.7 | *exsB* | Exoenzyme S synthesis protein B |
| PA14_42410 | –2.3 | | Hypothetical protein |
| PA14_42430 | –2.3 | *exsC* | Exoenzyme S synthesis protein C |
| PA14_42440 | –3.1 | *popD* | Translocator outer membrane protein PopD precursor |
| PA14_42450 | –3.5 | *popB* | Translocator protein PopB |
| PA14_42460 | –3.8 | *pcrH* | Regulatory protein PcrH |
| PA14_42470 | –3.8 | *pcrV* | Type III secretion protein PcrV |
| PA14_42480 | –4.1 | *pcrG* | Regulator in type III secretion |
| PA14_42520 | –3.6 | | Hypothetical protein |

| | | | |
|---|---|---|---|
| PA14_42540 | –2.3 | | Protein in type III secretion |
| PA14_42550 | –2.7 | *popN* | Type III secretion outer membrane protein PopN precursor |
| PA14_42570 | –2.4 | *pscN* | Type III secretion system ATPase |
| PA14_42580 | –3.8 | *pscO* | Translocation protein in type III secretion |
| PA14_42600 | –4.1 | *pscP* | Translocation protein in type III secretion |
| PA14_42610 | –4.5 | *pscQ* | Type III secretion system protein |
| PA14_42620 | –7.2 | *pscR* | Type III secretion system protein |
| PA14_42660 | –2.3 | *pscU* | Translocation protein in type III secretion |
| PA14_42950 | –2.0 | *fha2* | Type VI secretion system-associated FHA-domain protein TagH |
| PA14_42990 | –2.2 | *hsiH2* | Type VI secretion system baseplate subunit TssG |
| PA14_43000 | –2.5 | *hsiG2* | Type VI secretion system baseplate subunit TssF |
| PA14_43020 | –3.6 | *hsiF2* | Type VI secretion system baseplate subunit TssE |
| PA14_43030 | –3.2 | *hsiC2* | Type VI secretion system contractile sheath large subunit |
| PA14_43040 | –3.5 | *hsiB2* | Type VI secretion system contractile sheath small subunit |
| PA14_43050 | –3.1 | *hsiA2* | Type VI secretion system protein TssA |
| PA14_43070 | –2.4 | *hcp2* | Hcp family type VI secretion system effector |
| PA14_43090 | –2.4 | *tap* | DUF4123 domain-containing protein |
| PA14_43160 | –3.4 | | Transporter |
| PA14_45950 | –3.8 | *rsaL* | Regulatory protein RsaL |
| PA14_47160 | –2.3 | *cyoD* | Cytochrome *o* ubiquinol oxidase subunit IV |
| PA14_47180 | –2.6 | *cyoC* | Cytochrome *o* ubiquinol oxidase subunit III |
| PA14_47190 | –2.5 | *cyoB* | Cytochrome *o* ubiquinol oxidase subunit I |
| PA14_47210 | –2.7 | *cyoA* | Cytochrome *o* ubiquinol oxidase subunit II |
| PA14_48040 | –2.9 | *aprI* | Alkaline proteinase inhibitor AprI |
| PA14_48060 | –3.0 | *aprA* | Alkaline metalloproteinase |
| PA14_48090 | –3.0 | *aprF* | Alkaline protease secretion outer membrane protein AprF precursor |
| PA14_48100 | –2.7 | *aprE* | Alkaline protease secretion protein AprE |
| PA14_48115 | –2.6 | *aprD* | Alkaline protease secretion protein AprD |
| PA14_48140 | –2.4 | | Hypothetical protein |
| PA14_48530 | –2.4 | | AMP-binding protein |
| PA14_49230 | –2.2 | *napD* | NapD protein of periplasmic nitrate reductase |
| PA14_49260 | –2.1 | *napB* | Cytochrome *c*-type protein NapB precursor |
| PA14_49310 | –2.9 | | Hypothetical protein |
| PA14_49750 | –2.6 | | MFS family transporter |
| PA14_49760 | –2.4 | *rhlC* | Rhamnosyltransferase 2 |
| PA14_51350 | –4.1 | *phnB* | Anthranilate synthase component II |
| PA14_51360 | –5.5 | *phnA* | Anthranilate synthase component I |
| PA14_51380 | –5.4 | *pqsE* | Quinolone signal response protein |
| PA14_51390 | –5.6 | *pqsD* | 3-Oxoacyl-ACP synthase |
| PA14_51410 | –6.0 | *pqsC* | 3-Oxoacyl-ACP synthase III family protein |
| PA14_51420 | –6.2 | *pqsB* | Hypothetical protein |
| PA14_51430 | –5.9 | *pqsA* | AMP-binding protein |
| PA14_53250 | –3.2 | *cpbD* | Chitin-binding protein CbpD |
| PA14_53260 | –2.0 | | Hypothetical protein |
| PA14_59410 | –2.5 | | Hypothetical protein |

| | | | |
|---|---|---|---|
| PA14_59430 | –2.5 | | Hypothetical protein |
| PA14_59470 | –2.1 | | Hypothetical protein |
| PA14_59850 | –3.3 | | Hypothetical protein |
| PA14_59860 | –2.1 | | Type III effector Hop protein |
| PA14_59870 | –2.8 | | Hypothetical protein |
| PA14_60750 | –3.0 | *pra* | Protein activator |
| PA14_61870 | –2.3 | | Hypothetical protein |
| PA14_64060 | –2.5 | | Chemotaxis transducer |
| PA14_68430 | –2.5 | | Formate dehydrogenase accessory protein FdhD |
| PA14_68440 | –2.4 | | Oxidoreductase |
| PA14_68930 | –2.5 | | Permease |
| PA14_68940 | –3.1 | | Hypothetical protein |
| PA14_71900 | –5.2 | | Hypothetical protein |
| **Upregulated genes in PA14Δ32 compared to PA14** | | | |
| **ORF** | **$log_2$(fold-change)** | **Name** | **Product or function** |
| PA14_06790 | 2.1 | | Cytochrome *c* oxidase subunit |
| PA14_06800 | 2.3 | | Hypothetical protein |
| PA14_07850 | 2.3 | | ABC transporter substrate-binding protein |
| PA14_11660 | 3.0 | *aqpZ* | Aquaporin Z |
| PA14_11670 | 2.8 | | Hypothetical protein |
| PA14_13800 | 2.2 | *narH* | Respiratory nitrate reductase β subunit |
| PA14_13810 | 2.5 | *narJ* | Respiratory nitrate reductase δ chain |
| PA14_13830 | 2.2 | *narI* | Respiratory nitrate reductase γ chain |
| PA14_13840 | 2.1 | | Peptidyl-prolyl *cis-trans* isomerase/PpiC-type |
| PA14_13850 | 2.1 | *moaA* | Molybdenum cofactor biosynthesis protein A |
| PA14_21730 | 2.2 | | TonB-dependent receptor |
| PA14_26160 | 3.6 | | Hypothetical protein |
| PA14_26165 | 2.2 | | Hypothetical protein |
| PA14_28210 | 2.1 | | Hypothetical protein |
| PA14_28220 | 2.8 | | Hypothetical protein |
| PA14_28390 | 2.1 | | Hypothetical protein |
| PA14_31820 | 5.2 | | Aminotransferase |
| PA14_31840 | 5.0 | | Hypothetical protein |
| PA14_31850 | 5.1 | | Hypothetical protein |
| PA14_35320 | 3.4 | | 2-Hydroxyacid dehydrogenase |
| PA14_35330 | 3.2 | | 2-Ketogluconate transporter |
| PA14_35340 | 2.6 | | 2-Ketogluconate kinase |
| PA14_35360 | 2.4 | | Hypothetical protein |
| PA14_39650 | 2.1 | | TonB-dependent receptor |
| PA14_54690 | 3.2 | | Hypothetical protein |
| PA14_58580 | 2.2 | | Hydroxylase |
| PA14_61080 | 2.2 | | C4-dicarboxylate-binding protein |
| PA14_62260 | 2.3 | | Hypothetical protein |
| PA14_64790 | 3.5 | | MFS transporter |
| PA14_64800 | 5.9 | *vanA* | Vanillate *O*-demethylase oxygenase |
| PA14_64810 | 5.1 | *vanB* | Vanillate *O*-demethylase |

| | | | |
|---|---|---|---|
| PA14_72880 | 2.1 | | Short-chain dehydrogenase |
| PA14_72890 | 2.0 | | Transcriptional regulator |

**Table S7.** Expression of GCP genes in strain PA14Δ32.

| ORF | Name | $\log_2$(fold-change) |
|---|---|---|
| PA14_02110 | *siaD* | –1.3 |
| PA14_03720 | *pipA* | –0.7 |
| PA14_03790 | | –0.3 |
| PA14_04420 | | –0.6 |
| PA14_07500 | *rmcA* | –1.5 |
| PA14_16500 | *wspR* | –0.5 |
| PA14_20820 | *hsbD* | –0.4 |
| PA14_21190 | *nbdA* | –2.0 |
| PA14_21870 | | –0.8 |
| PA14_23130 | | –0.2 |
| PA14_26970 | | –0.8 |
| PA14_31330 | | –0.8 |
| PA14_37690 | | –0.4 |
| PA14_40570 | | –0.1 |
| PA14_42220 | *mucR* | –1.8 |
| PA14_45930 | *lapD* | –1.1 |
| PA14_49160 | | –0.4 |
| PA14_49890 | *yfiN/tpbB* | –0.4 |
| PA14_50060 | *roeA* | –0.5 |
| PA14_53140 | *rbdA* | –0.2 |
| PA14_53310 | | –0.6 |
| PA14_56280 | *sadC* | –1.5 |
| PA14_56790 | *bifA* | –0.8 |
| PA14_57140 | | –0.6 |
| PA14_60870 | *morA* | –1.2 |
| PA14_64050 | *gcbA* | –1.5 |
| PA14_65090 | *nicD* | –1.6 |
| PA14_65540 | *fimX* | –1.9 |
| PA14_66320 | *dipA* | –0.8 |
| PA14_69900 | *proE* | 0.0 |
| PA14_71850 | | 0.8 |
| PA14_72420 | *dgcH* | –0.6 |

**Table S8.** Minimal inhibitory concentration (MIC) for strains PA14 and PA14Δ32 exposed to different antibiotics.

| Antibiotic | MIC ($\mu$g mL$^{-1}$) using Sensititre™ plates | | |
|---|---|---|---|
| | PA14 | PA14Δ32 | PA14Δ32 + 23130[a] |
| Ampicilin | >16 | >16 | >16 |
| Ceftazidime | <2 | <2 | ≤4 |
| Ceftazidime/clavulanic acid | ≤1/4 | ≤1/4 | ≤1/4 |
| Cefotaxime/clavulanic acid | >2/4 | >2/4 | >2/4 |
| Cefotaxime | ≤16 | ≤16 | >32 |
| Ampicilin/sulbactam 2:1 ratio | >2/4 | >2//4 | >2/4 |
| Piperacilin/tazobactam constant 4 | <8/4 | <8/4 | <8/4 |
| Cefepime | ≤4 | ≤2 | ≤4 |
| Meropenem | <1 | ≤1 | <1 |
| Cephalothin | >32 | >32 | >32 |
| Imepenem | ≤1 | ≤1 | ≤1 |
| Amikacin | >8 | >8 | <8 |
| Levofloxacin | ≤0.25 | ≤0.5 | ≤0.5 |
| Gentamicin | <4 | <4 | >8 |
| Ciprofloxacin | ≤0.12 | ≤0.12 | ≤0.12 |
| Minocycline | >8 | >8 | ≤8 |
| Tigecycline | >2 | >2 | >2 |
| Cefuroxime | >16 | >16 | >16 |
| Ertapenem | >2 | ≤1 | ≤1 |
| Colistin | <1 | <1 | <1 |
| Cefoxitin | >16 | >16 | >16 |
| Doripenem | <4 | <4 | <4 |
| Rifampin | >8 | >8 | >8 |
| Nitrofurantoin | >64 | >64 | >64 |
| Fosfomycin (Fos) + glucose 6-phosphate | <Fos+ 32 | <Fos+ 32 | ≤Fos+64 |
| Trimethoprim/sulfamethoxazole | >32 | >32 | >32 |
| Chloramphenicol | >16 | >16 | >16 |
| Amoxicilin/clavulanic acid 2:1 ratio | >16/8 | >16/8 | >16/8 |
| Aztreonam | <16 | <16 | <16 |
| Nalidixic acid | >16 | >16 | >16 |

[a] Strain PA14Δ32 transformed with plasmid pJN_*23130*, carrying a wild-type copy of *PA14_23130*. Gentamicin resistance in this strain is connected to the presence of the expression plasmid.

**Table S9.** Phenotypic characterization of GCP-disrupted strains in different species.[a]

| Parameter | *Pseudomonas aeruginosa* | *Salmonella enterica* | *Caulobacter crescentus* | *Dickea zeae* | *Sinorhizobium* (*Ensifer*) *melilloti* |
|---|---|---|---|---|---|
| Number of GGDEF-domain proteins removed | 32 | 12 | 11 | 15 | 16 |
| Morphology | Affected | N.A. | Affected | N.A. | N.A. |
| Biofilm formation | Severely affected | Severely affected; cellulose synthesis requires c-di-GMP | Severely affected | Severely affected | Biofilm formation reduced but not abolished |
| Growth | Affected | N.A. | N.A. | N.D. | N.A. |
| Flagellum mediated motility | Swarming severely diminished; swimming not affected; normal flagella detected by TEM | Swarming and swimming severely affected; flagella lacking | Affected, flagella lacking; flagellar biogenesis and motility restored by c-di-GMP | Affected; swarming and swimming increase; higher number of flagella | N.A. |
| Pilli mediated motility (twitching) | Severely affected; type IV pili is not assembled and requires FimX | | Severely affected; type IV pili not assembled, assembly requires c-di-GMP | | |
| Virulence | Affected | Severely affected | | N.A. | N.A. (c-di-GMP not necessary for root nodule symbiosis) |
| Reference | This work | Solano et al. [100] | Abel et al. [101] | Chen et al. [102] | Schäper et al. [103] |

[a] *Abbreviations*: *N.A.*, not affected; *N.D.*, not determined.
